# Alternate routes to mnm^5^s^2^U synthesis in Gram-positive bacteria

**DOI:** 10.1101/2023.12.21.572861

**Authors:** Marshall Jaroch, Guangxin Sun, Ho-Ching Tiffany Tsui, Colbie Reed, Jinjing Sun, Marko Jörg, Malcolm E. Winkler, Kelly C. Rice, Troy A. Stich, Peter C. Dedon, Patricia C. Dos Santos, Valérie de Crécy-Lagard

## Abstract

The wobble bases of tRNAs that decode split codons are often heavily modified. In Bacteria tRNA^Glu, Gln, Asp^ contain a variety of xnm^5^s^2^U derivatives. The synthesis pathway for these modifications is complex and fully elucidated only in a handful of organisms, including the Gram-negative *Escherichia coli* K12 model. Despite the ubiquitous presence of mnm^5^s^2^U modification, genomic analysis shows the absence of *mnmC* orthologous genes, suggesting the occurrence of alternate biosynthetic schemes for the installation of this modification. Using a combination of comparative genomics and genetic studies, a member of the YtqA subgroup of the Radical Sam superfamily was found to be involved in the synthesis of mnm^5^s^2^U in both *Bacillus subtilis* and *Streptococcus mutans*. This protein, renamed MnmL, is encoded in an operon with the recently discovered MnmM methylase involved in the methylation of the pathway intermediate nm^5^s^2^U into mnm^5^s^2^U in *B. subtilis*. Analysis of tRNA modifications of both *S. mutans* and *Streptococcus pneumoniae* shows that growth conditions and genetic backgrounds influence the ratios of pathways intermediates in regulatory loops that are not yet understood. The MnmLM pathway is widespread along the bacterial tree, with some phyla, such as Bacilli, relying exclusively on these two enzymes. The occurrence of fusion proteins, alternate arrangements of biosynthetic components, and loss of biosynthetic branches provide examples of biosynthetic diversity to retain a conserved tRNA modification in nature.

**Importance:** The xnm^5^s^2^U modifications found in several tRNAs at the wobble base position are widespread in Bacteria where they have an important role in decoding efficiency and accuracy. This work identifies a novel enzyme (MnmL) that is a member of a subgroup of the very versatile Radical SAM superfamily and is involved in the synthesis of mnm^5^s^2^U in several Gram-positive bacteria, including human pathogens. This is another novel example of a non-orthologous displacement in the field of tRNA modification synthesis, showing how different solutions evolve to retain U34 tRNA modifications.

## Introduction

Post-transcriptional modifications of the first position of the anticodon of tRNA molecules (position 34, also called “wobble” position, Fig. 1A) are critical for accurate decoding of mRNAs (1). These modifications are often complex and can expand or decrease the wobble capacity that allows one given tRNA molecule to decode codons differing just by their third base (2). For example, bacterial tRNAs that decode NNA/NNG codons, particularly in split codon boxes, generally harbor xnm^5^(s^2^)U derivatives where nm^5^ stands for aminomethylene (3). The presence of the xnm^5^ side chain increases pairing to G-ending codons, while the s^2^ group may be the major contributor to restricting pairing with U and C-ending codons. In addition, some tRNAs carry ribose methylation at the 2’-OH and/or the mnm^5^ base modification, which may induce restrictive wobbling (4). The enzymes involved in introducing xnm^5^(s^2^)U derivatives are well-characterized in the model Gram-negative *E. coli* K12 (Fig. 1B) (see (5) for review). The insertions of the s^2^ and xnm^5^ moieties involve two separate biosynthetic branches, both requiring multiple enzymes (4, 5).

**Figure 1:**
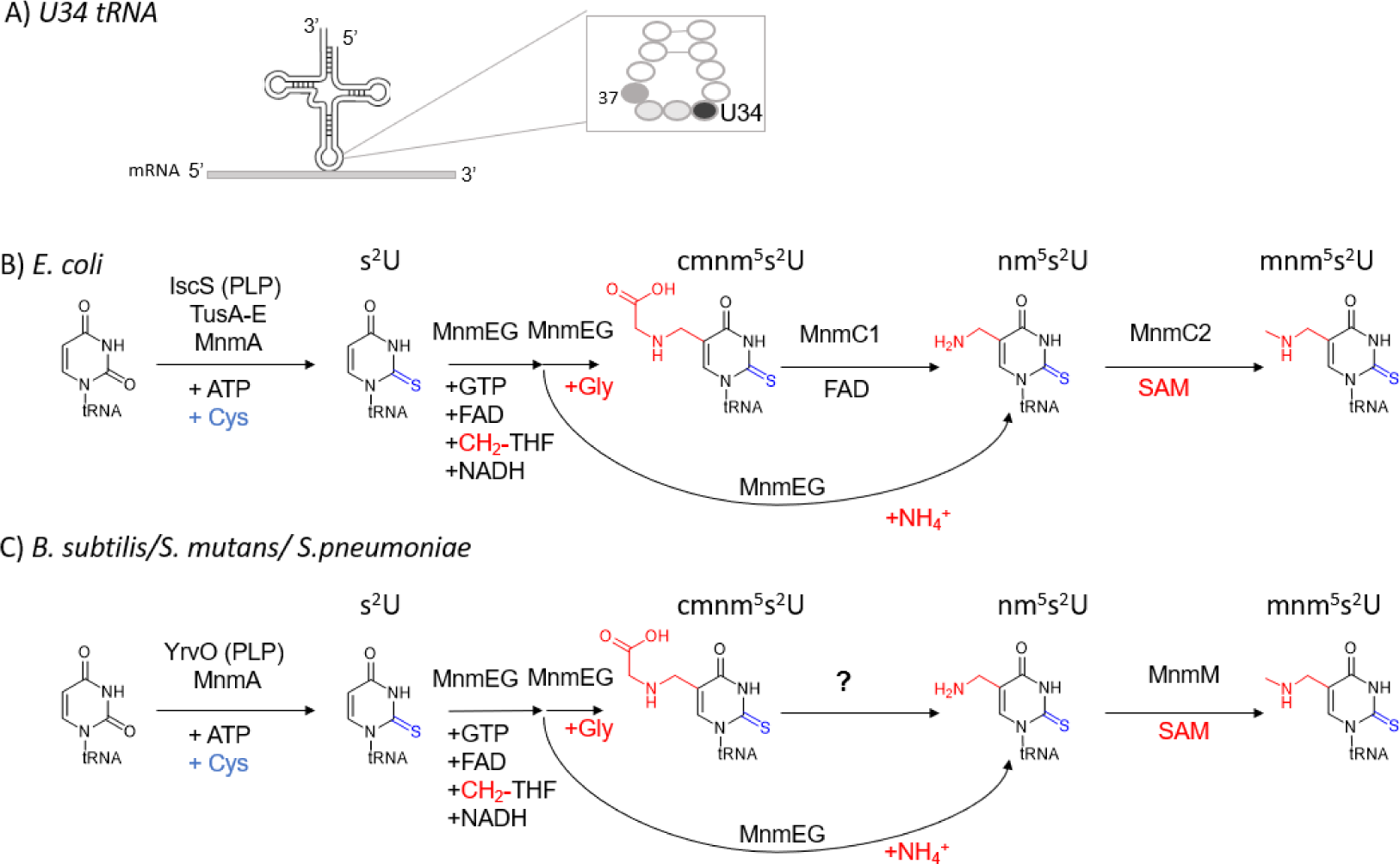
Validated and predicted xnm^5^(s^2^)U34 synthesis pathways in the two main model bacteria. (**A**) Wobble position of tRNAs. (**B**) Validated xnm^5^(s^2^)U34 synthesis enzymes for *E. coli* K12. (**C**) Validated xnm^5^(s^2^)U34 synthesis enzymes and pathway holes for *B. subtilis 168, S. mutans,* and *S. pneumoniae*. Abbreviations: 5-carboxymethylaminomethyl-2-thiouridine (cmnm^5^s^2^U); 5-carboxymethylaminomethyl-2’-O-methyluridine (cmnm^5^Um); 5-aminomethyl-2-thiouridine (nm^5^s^2^U); 5-methylaminomethyluridine (mnm^5^U); 5-methylaminomethyl-2-thiouridine(mnm^5^s^2^U).

The first step in the synthesis of the xnm^5^ moiety is catalyzed by a heterodimeric 5-carboxymethylaminomethyluridine-tRNA synthase complex MnmGE that can use glycine or ammonium as substrates to produce the cmnm^5^U or nm^5^U derivatives, respectively (6, 7) (Fig. 1B). The cmmn^5^U derivative can be subsequently converted into mnm^5^U by the bifunctional enzyme MnmC which contains two independent catalytic domains. The C-terminal domain of MnmC, also known as MnmC/MnmC1 or MnmC(o), promotes the FAD-dependent oxidoreduction of cmnm^5^s^2^U34 to form nm^5^U. Whereas the N-terminal MnmC domain, MnmD/MnmC2 or MnmC(m), harbors the tRNA (mnm^5^s^2^U34)-methyltransferase (EC:2.1.1.61 domain), which catalyzes the methylation of nm^5^s^2^U to nmn^5^s^2^U using SAM as the methyl donor (8)(Fig. 1B). Some organisms such as *Aquifex aeolicus* only encode a stand-alone MnmC2 protein and lack the MnmC1 ortholog as their MnmGE enzymes only use ammonium to directly produce nm^5^(s^2^)U (9). Other bacteria, such as *Mycoplasma capricolum,* only encode the MnmEG genes and no MnmC1 or MnmC2 genes and harbor cnmm^5^ (s^2^)U in their tRNAs (10).

Environmental and nutritional conditions affect the relative levels of tRNA modifications, including xnm^5^(s^2^)U derivatives (11), leading to varying translation efficiency and cellular viability. In general, media, growth phase, and tRNA identity influence the proportions of the different routes, leading to different xnm^5^(s^2^)U derivatives (12). It is also known that additional enzymes are involved in ribose methylation (TrmL)(13) or in producing tRNA^Lys^ and tRNA^Glu^ with U34 modified with selenium instead of sulfur (MnmH) [see (4) for review]. The MnmH enzyme is also involved in the geranylation of the same tRNAs in *E. coli* and *Salmonella typhimurium* (4). The presence of xnm^5^(s^2^)U modifications is critical for fitness as it has been shown in *E. coli* that mutations in *mnmE* and *mnmA* are synthetic lethal (14, 15). Mutations in *mnmE*, *mnmG,* and *mnmA* confer a wide range of phenotypes, including defects in virulence in diverse bacteria, including species from Gram-positive and Gram-negative groups (3, 16). Their orthologs are also important for modification of mitochondrial tRNAs and their absence leads to mitochondrial dysfunction and disease (3, 17).

*E. coli* K12 is one of the rare organisms where all the tRNA modification genes have been identified (18). Even in the well-studied Gram-positive model organism *Bacillus subtilis* sp subtilis strain 168, the functional assignment of genes involved in mnm^5^s^2^U synthesis remains incomplete (19). While the accumulation of cmnm^5^sU was anticipated, given the presence of phylogenetically conserved *mnmEG* orthologs in this organism, the presence of mnm^5^s^2^U was initially unexpected since *B. subtilis* lacks *mnmC*-like gene or sequences coding for MnmC1 and MnmC2 domains. This intriguing finding suggested that a non-orthologous replacement must have occurred for the last steps of the pathway (Fig. 1C) (20). The existence of alternate biosynthetic schemes for tRNA thiolation in this organism is further supported through the occurrence of an abbreviated pathway for s^2^U tRNA modification involving only two enzymes YrvO-MnmA, bypassing the Tus sulfur relay system required in *E. coli* (Fig. 1C) (21).

Foundational studies from the Armengod group showed that cell extracts of *B. subtilis* were competent to promote mnm^5^s^2^U synthesis from precursor cmnm^5^s^2^U tRNA isolated from *E. coli ΔmnmC* strain) (20). Interestingly, these reactions also formed the nm^5^s^2^U intermediate, which was consumed in reactions supplemented with S-adenosyl-methionine (SAM), but not FAD. More recently, YtqB (MnmM) was identified as the enzyme responsible for the conversion of nm^5^s^2^U into mnm^5^s^2^U, performing an MnmC2-like reaction (22). This assignment was attributed to the depletion of mnm^5^s^2^U accompanied by the accumulation of nm^5^s^2^U tRNA in *B. subtilis ΔmnmM* strain. Trans-complementation of not only *B. subtilis mnmM*, but also *Staphylococcus aureus* and *Arabidopsis thaliana mnmM* orthologs restored mnm^5^s^2^U levels (22). Furthermore, according to Cho and collaborators, *B. subtilis mnmM* expression in *E. coli ΔmnmC* shows detectable levels of mnm^5^s^2^U but at levels significantly lower than those of the wild type. While MnmM is competent in catalyzing methylation of nm^5^s^2^U to yield mnm^5^s^2^U, it remains unanswered if additional components participate in this biosynthetic scheme in *B. subtilis* and other species lacking MnmC. For instance, although experimentally demonstrated in crude extracts, the identity of an MnmC1-like enzyme catalyzing the conversion of cmnm^5^s^2^U into nm^5^s^2^U tRNA is unknown.

Comparative genomic methods combining phylogenetic distribution, gene fusion, and physical clustering analyses (21) have been powerful tools in identifying many missing tRNA modification genes, particularly for complex modification pathways that require many enzymes (23–27). Now that many tRNA modification genes have been identified in different organisms, it is clear that different evolutionary solutions have emerged to catalyze the insertion of the same modifications (28). In this work, we used comparative genomic methods combined with genetic studies to better understand the last steps in nm^5^s^2^U synthesis in microbes lacking MnmC and identified the versatile radical SAM superfamily family protein YtqA as a key player. These combined approaches reveal the occurrence of alternate pathways leading to hypermodification of U34 tRNA^Glu, Gln, Lys^ in bacterial species.

## Results

### Phylogenetic distribution analysis of mnm^5^s^2^U pathway suggests MnmC1 has been replaced by non-orthologous enzymes in a number of firmicutes

Analysis of the distribution of the known mnm^5^s^2^U34 synthesis protein family members in the bacterial genomes covered in the COG database revealed a patchy distribution for the latter part of the pathway (Fig. 2, Fig. S1, Supplemental data S1 and S2). Of the 968 organisms that encode the three MnmA (COG0482/K00566), MnmE (COG0486/K03650), and MnmG (COG0445/K03495) proteins, 22% (216/968) encode bifunctional MnmC1/MnmC2 (or MnmC, COG4121/K15461) proteins like *E. coli,* while ∼24% (232/968) only encode only MnmC2 proteins without MnmC1 domains like *A. aoelicus*. Finally, over half (520/968) of the analyzed organisms lack both MnmC1 and MnmC2 homologs. Thus, the occurrence of MnmC1 is sparse and matches the distribution of MnmC1 and MnmC2 sequences recently reported by Cho et al. (22) for a smaller dataset. The absence of MmnC1 can be, however, associated with distinct biochemical scenarios. First, in the subset of organisms that only use the ammonia pathway to directly synthesize nm^5^s^2^U, no MnmC1 is required, and the subsequent methylation to mnm^5^s^2^U can be performed by MnmC2 family proteins, like in *A. aeolicus* (9) or possibly by YtqB/MnmM proteins (22). Second, some organisms such as *Lactococcus lactis* sub lactis Il1403 (29) or *M. capricolum* (10) harbor only cmnm^5^s^2^U and, therefore, do not require any enzymes beyond MnmEG. Third, in species that accumulate both cmnm^5^s^2^U and mnm^5^s^2^U, as in the case of *B. subtilis* and possibly in many other Firmicutes (30, 31) (Fig. 2 and Fig. S1), a non-orthologous replacement of MnmC1 must be present and is yet to be identified.

**Figure 2.**
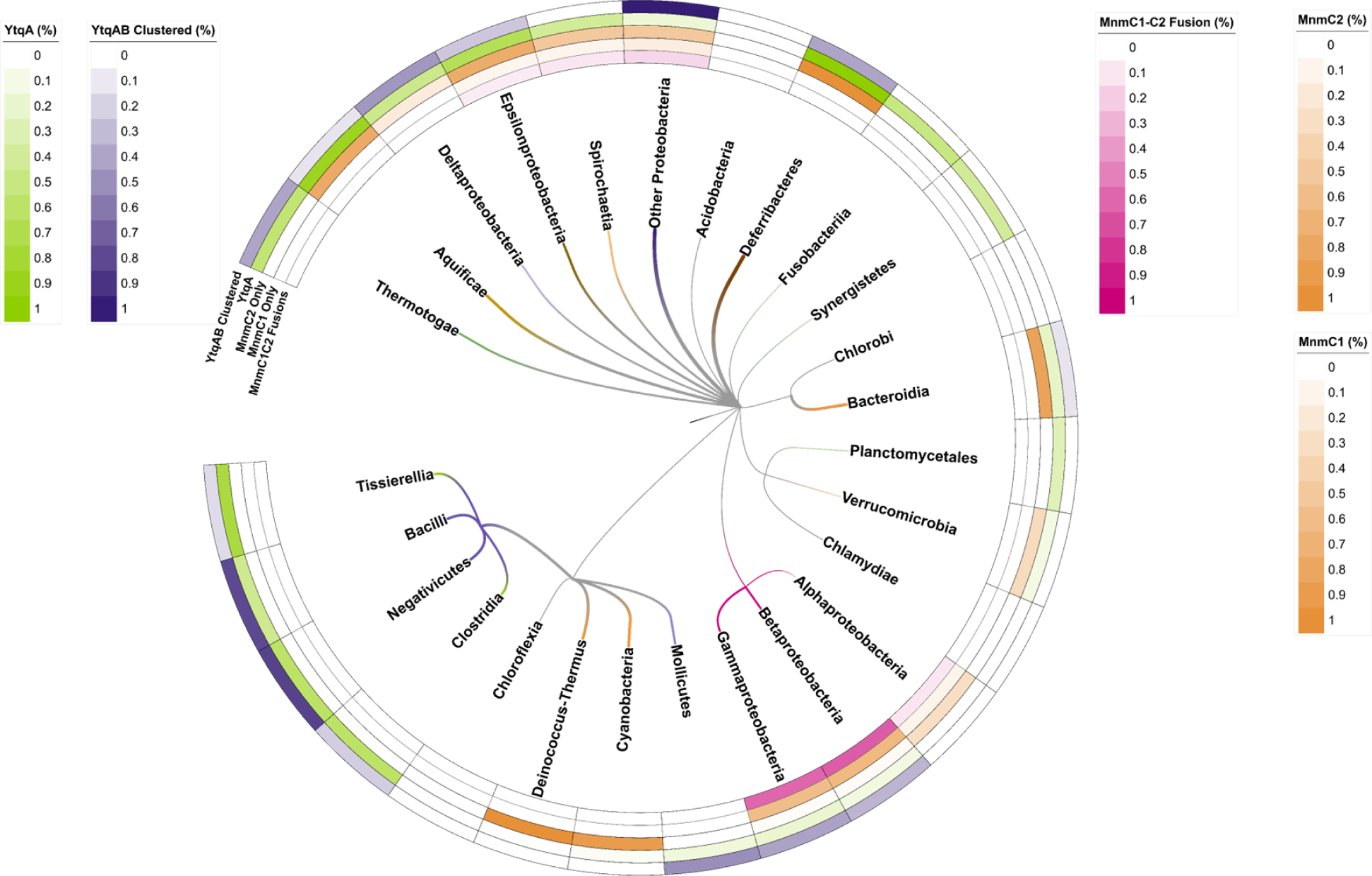
Phylogenetic distribution analysis of MnmC1, MnmC2, and YtqA/YhcC proteins. Percentages for each group represent the number of organisms that encode a given protein group [MnmC1/MnmC2 fusions (MmmC1C2), MnmC1 alone (MnmC1), MnmC2 alone (MnmC2), and YtqA/YhcC] out of the 968 bacterial genomes that encode MmnAGE analyzed (Table S1; Supplemental data S2, S3, and S4). In addition, the percentage of genomes where the *ytqA* and *mnmM* genes are physically clustered on the chromosome (YtqA-MnmM clustered) is reported (Supplemental Data S5 and S6). The pixel width of branches reflects the relative intensity of corresponding heatmaps and their dominant (highest value) color of those same heatmaps (0.0-1.0 heatmap range corresponds to 1-3 px branch width). Heatmap per branch with the highest intensity across all five heatmap rows was assigned to the branch color to highlight taxonomic patterns. Only Deferribacteres possess a combined color reflecting the paired 1.0 values for both MnmC2 and YtqA/YhcC heatmaps (orange and green of equal intensity combine to make brown). The Firmicute branch is colored red.

### Comparative genomic analyses led to the identification of a Radical SAM family subgroup as a candidate for the missing mnm^5^s^2^U synthesis enzyme in Firmicutes

Protein fusions and physical clustering analysis are powerful methods to identify missing genes (32, 33) and both led to the identification of a subgroup of the YtqA/YhcC family as a strong candidate for an enzyme involved in the synthesis of nm^5^s^2^U in Firmicutes. A domain analysis of the COG4121 family, which encompasses both MnmC1 and MnmC2 domains, across the 968 bacterial genomes of the COG Database (34) that also encode the three MnmA, MnmG, and MnmE proteins, led to the identification of at least four other types of MnmC2 fusions, in addition to the well-characterized MnmC1/MnmC2 examples: fusions with the queuine tRNA-ribosyltransferase (TGT) (COG0343, PF01702); the DNA modification methylase YhdJ (PF01555, COG0863); the menaquinone biosynthesis MqnA superfamily subgroup MqnD (PF02621, cd13634); and with a radical SAM family domain of the YtqA/YhcC subgroup (COG1242, PF16199, cd01335, PF04055) (Supplemental data S3, Figure S1). The YtqA name comes from the subfamily member found in *B. subtilis* (BSU30480) and the YhcC name comes from the subfamily member found in *E. coli* (b3211). Three examples of YtqA/MnmC2 fusion proteins were identified in the genomes present in the COG database and 70 more in the InterPro database (Fig. 3A, Figure S1, Supplemental data S4A). Genomic synteny analyses show that *ytqA* is often in an operon or in a close physical neighborhood with the *mnmM* gene encoding the recently identified nm^5^s^2^U methyltransferase, including in the model Gram-positive *B. subtilis* where the two genes are co-expressed (Fig. 3B, Fig. 4B and Fig. S2) (35). Both YtqA/YhcC and MnmM are part of superfamilies that are currently not well separated in most protein family databases (see below and discussion), but when analyzing the distribution of *ytqA-mnmM* clusters (Fig. 2 and Supplemental data 5), these are prevalent in organisms that are missing MnmC1 and MnmC2 such as the Firmicutes and the deltaproteobacteria (Fig. 2, Supplemental data S6, data S7).

**Figure 3.**
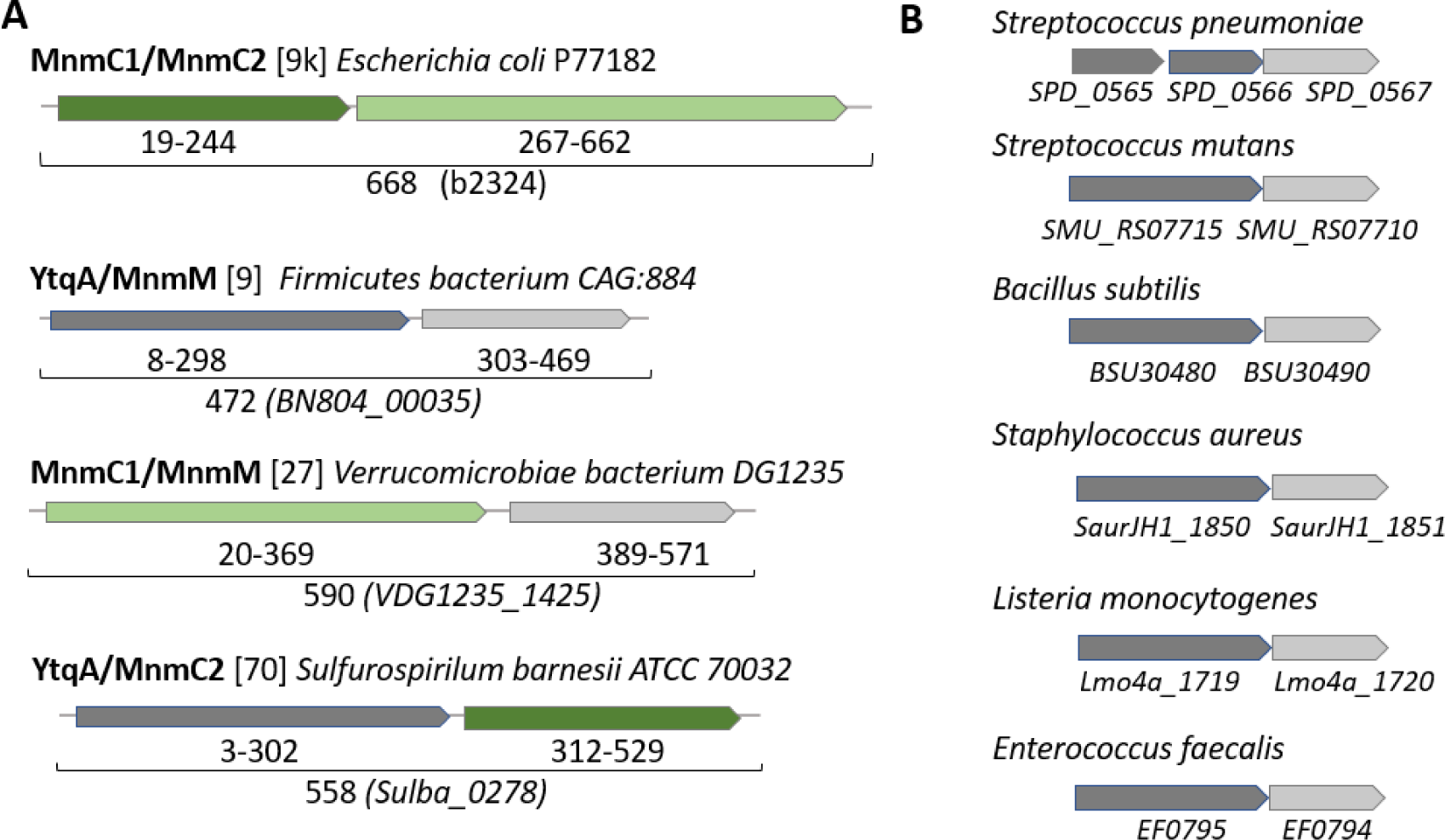
Physical clustering and domain fusion link the YtqA subgroup with mnm^5^s^2^U synthesis. (**A**) Examples of fusions between proteins involved in the last steps of mnm^5^s^2^U synthesis. Protein domains are color-coded MnmC1 (dark green), MnmC2 (light green), MnmM (light gray), and YtqA (dark gray). The number of examples of such fusions present in the InterPro database is given between brackets; see supplemental data S4A for the complete list of YtqA/MnmM, MnmC1/MnmM, and YtqA/MnmC2 fusions. (**B**) Examples of physically clustered *ytqA* (dark gray) and *mnmM* (light gray) in Gram-positive species. Data extracted from the SSN analysis performed for Fig. 4.

**Figure 4.**
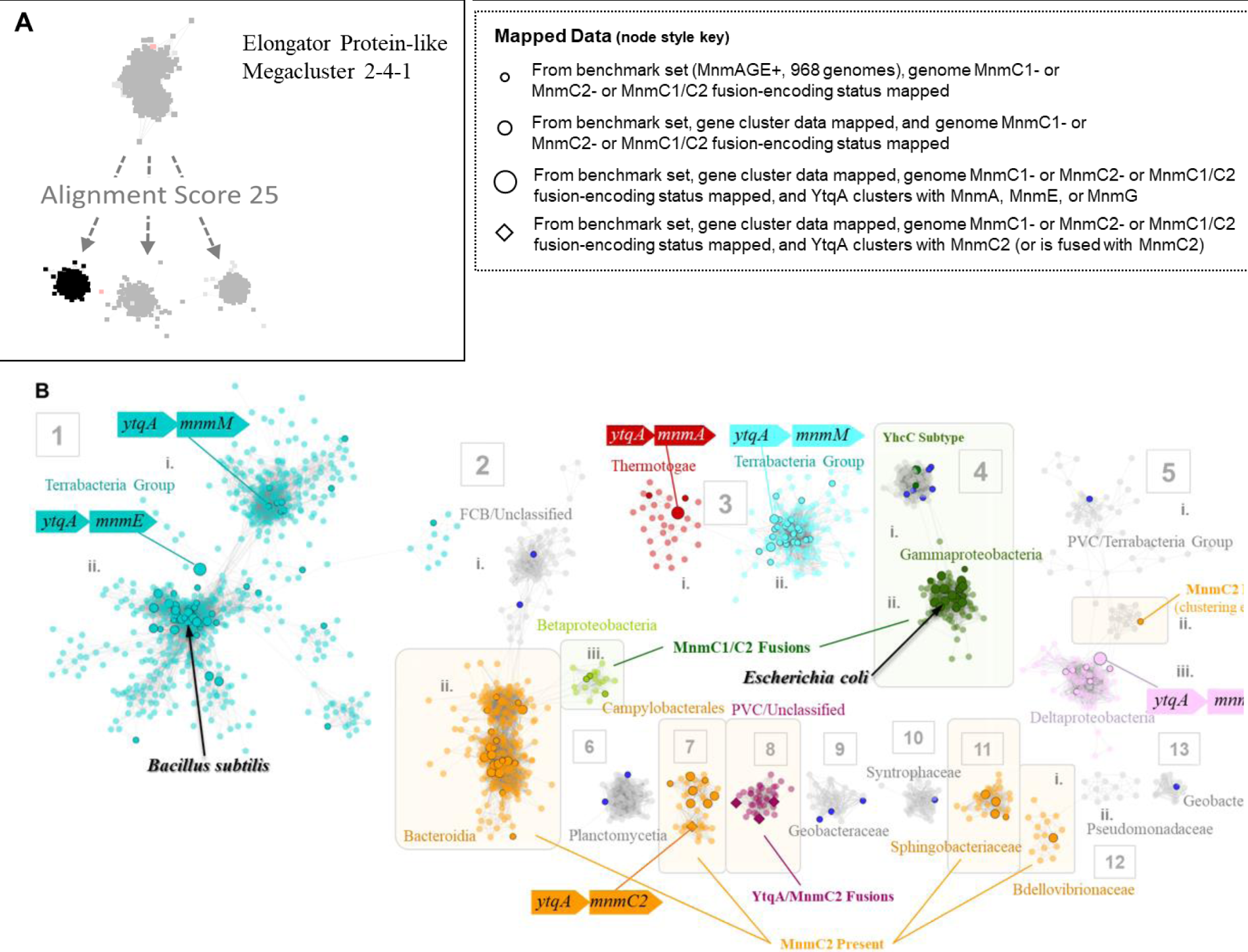
Sequence similarity network of the YtqA/YhcC family. (**A**) The radical SAM domains that are fused to MnmC2 are members of the Megacluster-2-4-1 (Elongator protein-like) on RadicalSAM.org (in black), a subgroup of Megacluster 2.4; (**B**) YtqA/YhcC SSN overview generated using InterPro HMM signature identifier IPR005911 (EFI-EST; AS100, repnode 55). Different types of data were mapped to the subset of nodes (with black outlines) present in the benchmark genomes (968 representative genomes that encode MnmAGE proteins: 1) Presence/absence data of MnmC1, MnmC2 and MnmC1/C2 fusions (Supplemental data S3); YtqA/MnmC2 fusions data (Supplemental data S3); physical clustering data (Supplemental data S5). Dark blue nodes were found to be without Gene Cluster data in KEGG. For complete mapping, see Supplemental Data S8.

YhcC/YtqA sequences display signature motifs of members of the radical SAM superfamily of enzymes, known to facilitate a wide range of chemistries. First, these sequences were located in the sequence similarity network (SSN) analysis of the whole radical SAM family generated by the Gerlt group using the searchable web resource (RadicalSAM.org)(36). The YtqA/YhcC subgroup was identified as part of the Elongator Protein-like mega cluster 2-4.1 (Fig. 4A). Proteins of this subgroup are members of the KEGG orthology group K07139 and the InterPro family IPR005911 (Protein YhcC-like). YtqA unique functional motifs and genomic synteny of its coding sequence make it a candidate for replacing MnmC1 in many organisms. Proteins within the YhcC sub-group would not be isofunctional since E coli genome encodes for YhcC, and MnmC1 is the only protein catalyzing the cmnm^5^s^2^U to nm^5^s^2^U reaction in this organism (8). We, therefore, constructed a SSNs of 10,896 IPR005911 proteins, which allowed differentiation of the *E. coli* YhcC subgroup (group 4 in Fig. 4B) from the *B. subtilis* YtqA subgroup (group 1 in Fig. 4B). Mapping of additional data such as gene neighborhood, fusion and gene presence/absence data onto this network suggests that in addition to YtqA proteins from subgroup 1, homologs from subgroups 3, 5, 7 and 8 are most likely linked to mnm^5^s^2^U synthesis. Collectively, bioinformatic analysis showing the physical clustering, gene fusion, and taxonomic distribution data strongly suggest a role of the YtqA protein subfamily in Firmicutes in mnm^5^s^2^U synthesis in tRNA.

### Genetic analysis in *S. mutans* confirms the involvement of YtqA in mnm^5^s^2^U synthesis

To validate the predicted role of YtqA in mnm^5^s^2^U synthesis, we constructed deletion mutants of the *ytqA and mnmM genes* in *Streptococcus mutans*. Like in *B. subtilis*, the two genes are in an operon in *S. mutans (*Fig. 3). A non-polar deletion of the *ytqA* (*smu_1699c*) was constructed by inserting an Erythromycin Resistance (Ery^R^) cassette lacking a terminator. The same cassette was used to delete the *mnmM* (*ytqB*, *smu_1697c*) gene as well as to delete both genes. As expected, tRNA extracted from a *Streptococcus mutans* strain containing a deletion of the *ytqA-mnmM* genes (*smu_1699c and smu_1697c)* lacks mnm^5^s^2^U and accumulates increased relative levels of nm^5^s^2^U and cmnm^5^s^2^U intermediates (Fig. 5). Inactivation of only *mnmM* confirms the involvement of the methyltransferase in mnm^5^s^2^U formation and also showed an accumulation of nm^5^s^2^U, compatible with its proposed role in catalyzing the final methylation step in mnm^5^s^2^U. Inactivation of *ytqA* led to lower levels of mnm^5^s^2^U and the loss of nm^5^U accumulation, a phenotype that supports the involvement of *ytqA* in this pathway. Notably, the distinct phenotypes of *ytqA-mnmM* and *ytqA* deletions demonstrate the absence of polar effects on *ytqA* knockout strain.

**Figure 5.**
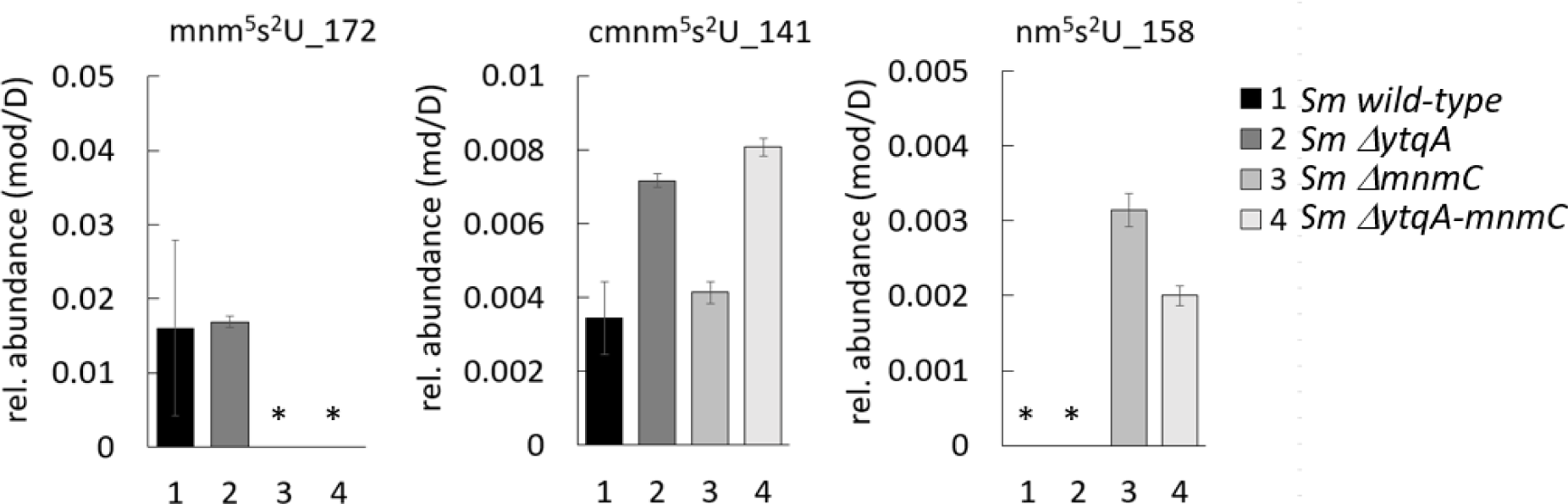
*S. mutans* YtqA and MnmM are involved in mnm^5^s^2^U tRNA modification. *S. mutans* wild-type, and *ytqA*, *mnmM,* and *ytqAmnmM* deletion strains were cultured in BHI medium. tRNA samples were processed through the VDCL2 method and analyzed through LC-MS method 2, as described in the methods section. For each sample, 600 ng of digested tRNA was analyzed through LC-MS, and two technical replicates were analyzed. The reported averages and standard deviation were calculated based on data acquired from three independent growth experiments. The signal of each analyte was confirmed by both qualifier and quantifier transitions and reported as the fragment mass of each modification, as indicated. Relative levels of each modification were determined by normalizing the area of mass abundance associated with each modification to the mass abundance of dihydrouridine in the same sample. The star mark denotes the absence of modification or levels below the detection limit.

### Genetic analysis of *B. subtilis* YtqA shows its involvement in mnm^5^s^2^U pathway

The genomic co-localization of *ytqA* and *mnmM* observed in *S. mutans* is also observed in *B. subtilis*. Recently, MnmM has been identified as the SAM-dependent methyltransferase catalyzing the final step in mnm^5^s^2^U synthesis (22) In this report, Cho et showed that *nmnM* inactivation resulted in the loss of mnm^5^s^2^U with concomitant accumulation of nm^5^sU, a phenotype similar to that of *S. mutans*. However, earlier studies from the Armengod laboratory had shown that inactivation of *mnmM* (*ytqB*) does not eliminate the synthesis of mnm^5^s^2^U tRNA (20). To clarify these opposing findings, we analyzed the relative levels of tRNA modifications in *B. subtilis ytqA* and *mnmM* mutants independently in two different laboratories. These analyses performed by two research groups showed the absence of mnm^5^s^2^U modification in the Δ*ytqA* strain and reduced levels in the *ΔmnmM* strain from LB cultures isolated at mid and late lag phase (OD 0.5-1) (Fig. 6 and S3A). In our hands, tRNA samples extracted from *B. subtilis* cultured beyond log phase (OD = or > 2) led to inconsistent results (Fig. S3B), likely associated with cells entering sporulation. Remarkedly, tRNA isolated from *B. subtilis ΔytqA* showed high levels of the pathway intermediate nm^5^s^2^U that is not detectable in tRNA extracted from the wild-type strain. These results are in contrast with the recent Cho et al report on the analysis of *mnmM* knockout strain using tRNA samples isolated from overnight cultures that showed no detectable levels of mnm^5^s^2^U (22), but in agreement with earlier analysis of *B. subtilis* IC6779 strain containing *mnmM* inactivation (20). Overall, these analyses showed the involvement of both genes in mnm^5^s^2^U. More specifically, *ytqA* inactivation resulted in the accumulation of nm^5^s^2^U and no detection of mnm^5^s^2^U. It is worth noting that cells lacking YtqA or MnmM do not display any growth phenotype when compared to the wild type, suggesting that the intermediate cmnm^5^s^2^U modification may already provide the proper conformation of s^2^U34 to guarantee fidelity in base pairing with cognate tRNAs.

**Figure 6.**
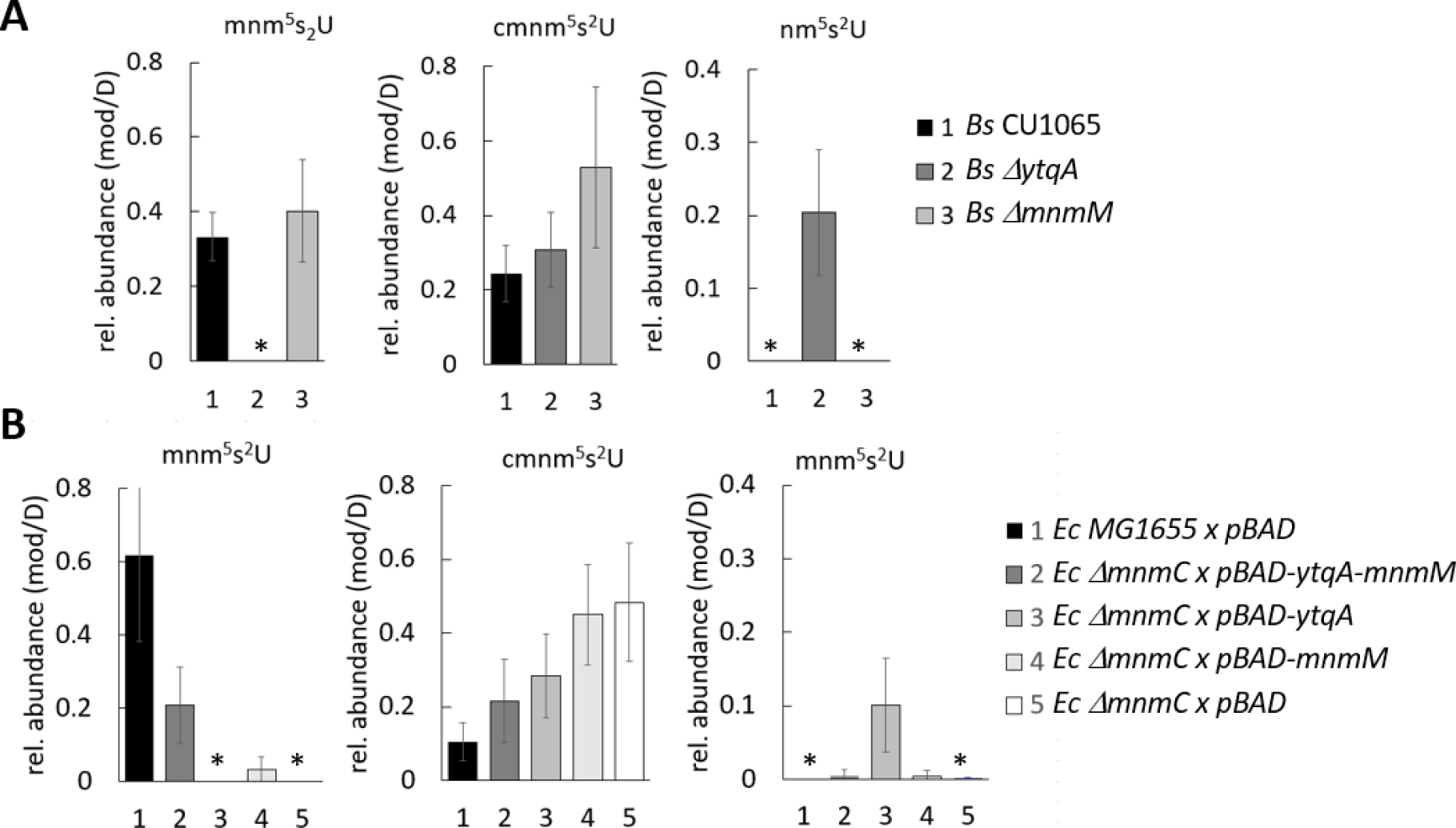
*B. subtilis* mnm^5^s^2^U pathway involves YtqA and MnmM. **A)** Inactivation of *ytqA* and *mnnM* genes affects the relative accumulation of mnm^5^s^2^U and pathway intermediates. Modified nucleosides were extracted from *B. subtilis* wild type, *ΔytqA* and *ΔmnmM* strains were cultured in LB medium until OD of 1. **B)** *B. subtilis* YtqA-MnmM complements defects associated with *E. coli* Δ*mnmC* inactivation*. E. coli* wild type MG1655 *and* Δ*mnmC* strain transformed cells with empty plasmid (pBAD), pDS358 (*pBAD-ytqA-mnmM*), pDS359 (*pBAD-ytqA*), or pDS365 (*pBAD-mnmM*) were cultured in LB ampicillin and arabinose until OD of 1. tRNA samples were isolated using the PDS method and analyzed using LC-MS method 1. Relative levels of each modification were determined by normalizing the mass abundance associated with each modification to the mass abundance of dihydrouridine in the same sample. The reported averages were calculated based on data acquired from three independent growth experiments. The star mark denotes the absence of modification or levels below the detection limit.

The ability of *B. subtilis* YtqA and MnmM to functionally replace *E. coli* MnmC was then assessed through heterologous trans-complementation experiments as a probe for MnmC1 and MnmC2 equivalent reactions. As previously reported, *E. coli ΔmnmC* accumulates cmn^5^s^2^U and completely lacks mnm^5^s^2^U (20). Co-expression of *B. subtilis* YtqA-MnmM restored the synthesis of mnm^5^s^2^U to 1/3 of wild-type levels, confirming their involvement in this pathway (Fig. 6B). Expression of *B. subtilis* MnmM alone also resulted in mnm^5^s^2^U formation, albeit at much lower levels (∼5% of E. coli MG1655 levels) and similar to residual levels reported by Cho et al. (22).

### Altered pathway for the mnm^5^s^2^U synthesis in *S. pneumoniae*

Genomic analysis of YtqA and MnmM orthologs showed the presence of an altered pathway for mnm^5^s^2^U in some Gram-positive such as *Streptococcus pneumoniae* D39. This organism harbors truncated copies of *ytqA* coding the N-terminal and C-terminal sequences of YtqA *(spd_0565* and *spd_0566*) upstream of *mnmM (spd_0567*) (Fig. 3 and Fig. S4), raising the question of whether these species represent evolutionary representatives in the process of losing the last step of this pathway. In fact, analysis of modified nucleosides from *S. pneumoniae* showed much-reduced levels of mnm^5^s^2^U (approximately 10-fold lower than *B. subtilis* or *E. coli*, Fig. 6) and confirmed the involvement of *spd_565* to *spd_567* gene region in this pathway (Fig. 7). Interestingly, a spontaneous nonsense mutation within *mnmM* generating a Q152stop was identified within the unencapsulated *S. pneumoniae* D39 genetic background (IU1824) (Fig. S4). Analysis of modified nucleosides showed that this truncation near the C-terminal end of the protein affects MnmM activity and results in loss of mnm^5^s^2^U synthesis like the deletion of the whole *mnmM* gene (IU19966) (Fig. 7). These phenotypes were complemented by repairing the mutation (strain IU19835) or expressing the *mnmM* ectopically under a zinc inducible promoter. The lack of an apparent growth defect upon elimination of mnm^5^s^2^U was also observed in *S. pneumoniae,* reinforcing the proposal that the cmnm^5^s^2^U modification is sufficient to fulfill functional requirements. Nevertheless, out of 242 complete *S. pneumoniae* genomes in the BV-BRC database at the time of the analysis (version 3.32.13a), all harbored truncated copies of *ytqA (e*xamples given in Fig. S5), suggesting this truncation must have occurred just before the *S. pneumoniae* speciation. Phylogenetic analysis indicates that the closest species with a common ancestor, *Streptococcus mitis* (37) has a full copy of the gene (Fig. S5).

**Figure 7.**
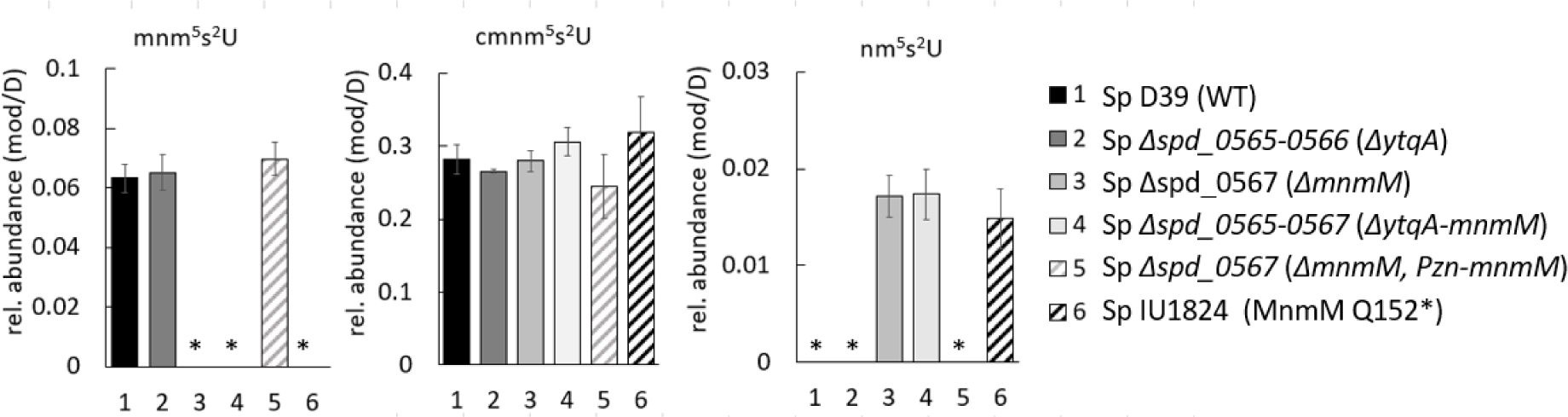
*S. pneumoniae* (Sp) MnmM is involved in mnm^5^s^2^U tRNA modification. tRNA isolated from Sp cultures using MW methods were digested and analyzed through LC-MS Method 1. Modified nucleosides were purified tRNA from Sp strains: 1) Sp IU19835: D39 Δ*cps* WT (*spd_0565^+^*, *spd_0566^+^, spd_0567^+^*) (black), 2) Sp IU19964: Δ*spd_0565-0566* markerless (*ΔytqA*) (dark grey), 3) Sp IU19966: *Δspd_0567* markerless (*ΔmnmM*) (light grey),4) Sp IU19968: Δ*spd_0565-0567* markerless (*ΔytqAmnmM*)(white), 5) Sp IU19974: Δ*spd_0567* markerless (*ΔmnmM*) ΔbgaA-Pzn-*mnmM* grown in Zn, (complementation of *ΔmnmM*) (grey stripped), and 6) Sp IU1824: original WT strain (IU1824) that has *spd_0567* (Q152stop) mutation (black stripped). *spd_0565* and *spd_0566* encode the N-terminal and C-terminal of YtqA and *spd_0567* encodes MnmM. Relative levels of each modification were determined by normalizing the mass abundance associated with each modification to the mass abundance of dihydrouridine in the same sample. The reported averages were calculated based on data acquired from three independent growth experiments. The star mark denotes the absence of modification or levels below the detection limit.

## Discussion

Bioinformatic analysis shows the occurrence of diverse biochemical strategies for hypermodification of U34 in tRNA^Glu, Gln, Lys^ in bacteria. Our results from the *S. mutans* and *S. pneumoniae ΔmnmM* strains confirm the functional assignment of Cho et al. (22) that MnmM is required in the SAM-dependent methylation reaction nm^5^s^2^U to form mnm^5^s^2^U. Our analysis from *B. subtilis ΔmnmM,* however, did not show the same requirement, suggesting that, in this organism, other methylases may provide functional replacement under certain growth conditions. Previous analysis of single knockout strains of several methylases in *B. subtilis*, including MnmM also failed to show inactivation of mnm^5^s^2^U synthesis (20). We believe that the differences in the observed phenotypes of the *mnmM* knockout strain between ours and other studies may be associated with the functionality of the NH_4_ and glycine-dependent routes with varying growth conditions and stages. The NH_4_ route could be the major route to mnm^5^s^2^U in the stationary phase (conditions used in the Cho et al. analyses). Hence MnmM would be the major player in these conditions. However, the glycine route may be the major route in exponential growth and another enzyme or pathway would be required to synthesize mnm^5^s^2^U from cmnm^5^s^2^U. Physical clustering and gene fusion data point to YtqA as a candidate to complete this pathway in *B. subtilis*. Indeed, the *B. subtilis ΔytqA* mutant is devoid of mnm^5^s^2^U and accumulates nm^5^s^2^U. This phenotype cannot be due to a polar effect on *mnmM,* as we found that deleting this gene did not lead to nm^5^s^2^U accumulation and retained mnm^5^s^2^U modification. In addition, co-expression of *ytqA-mmnM* allowed partial complementation of the *E. coli mnmC* mnm^5^s^2^U deficiency, whereas expression of *mmM* alone only led to minimal mnm^5^s^2^U accumulation. The expression of *ytqA* alone did not increase nm^5^U nor mnm^5^U levels in the same *mnmC* strain compatible with the model that thiolation of C2 of U34 precedes the installation of C5 modification route (data not shown).

The exact role of YtqA is still not fully understood and will require further biochemical and genetic characterization. Nevertheless, experimental evidence provided in this study shows the involvement of YtqA in the biosynthesis of nmn^5^s^2^U; therefore, we propose renaming this enzyme as MnmL (Figure 8). Our current model is that MnmL-MnmM can promote the conversion of cmnm^5^s^2^U into mnm^5^s^2^U. MnmL (YtqA) is classified as a member of radical SAM enzymes, which are known to catalyze a range of diverse chemical transformations triggered by the generation of a 5’-deoxyadenosyl radical (dAdo^●^) from SAM (38). This dAdo^●^ could abstract an H-atom from cmnm^5^s^2^U, causing oxidative decarboxylation by forming an xnm^5^s^2^U intermediate that could further react to yield mnm^5^s^2^U. Potential sites for the dAdo^●^-catalyzed H-atom abstraction include the glycine alpha carbon, the amide nitrogen, or the ^5^U-methylene carbon (Figure S6). Experimental determination of the site of H-atom abstraction would require isotope labeling studies. However, the viability of this chemistry can be probed using density functional theory (DFT) geometry optimizations to predict the outcome of H-atom abstraction from each of these sites within the cmnm^5^s^2^U substrate—these results are summarized in Table S2 and Figure S7. Briefly, H-atom abstraction from the amide nitrogen promotes oxidative decarboxylation and the formation of a terminal imine that could be hydrolyzed to give nm^5^s^2^U. This chemistry is very similar to that catalyzed by radical SAM enzymes HydG and ThiH, which abstract hydrogen from the amino group of an exogenous L-tyrosine, leading to the formation of *p*-cresol and dehydroglycine (39). H-atom abstraction from either of the other sites on cmnm^5^s^2^U potentially creates scissile intermediates that could lead to nm^5^s^2^U (Figure S7). However, the observation made in the present work that both YtqA and MnmM are required to produce the maximal amount of mnm^5^s^2^U suggests some interplay between the two enzymes. Sequence homology (cf. Figure 4) suggests that MnmL (YtqA) possesses a (β/α)_6_ three-quarter TIM barrel fold, like 75% of all radical SAM enzymes (40). Often, an additional protein subunit occupies the gap in the TIM barrel and is integral in the catalyzed chemistry, as is the case in the tRNA-modifying MiaB (41). MnmL is a quite small radical SAM enzyme (sequence length ≈330 amino acids), and no additional subunit is predicted by AlphaFold {https://alphafold.ebi.ac.uk/entry/O35008}. Therefore, it is possible that tRNA binding may mediate the formation of a ternary complex with MnmM, which enables the conversion of cmnm^5^s^2^U to mnm^5^s^2^U. This model is compatible with the observation that no nm^5^s^2^U is accumulated in *S. mutans*, *B. subtilis*, and *S. pneumoniae* wild-type strains.

**Figure 8:**
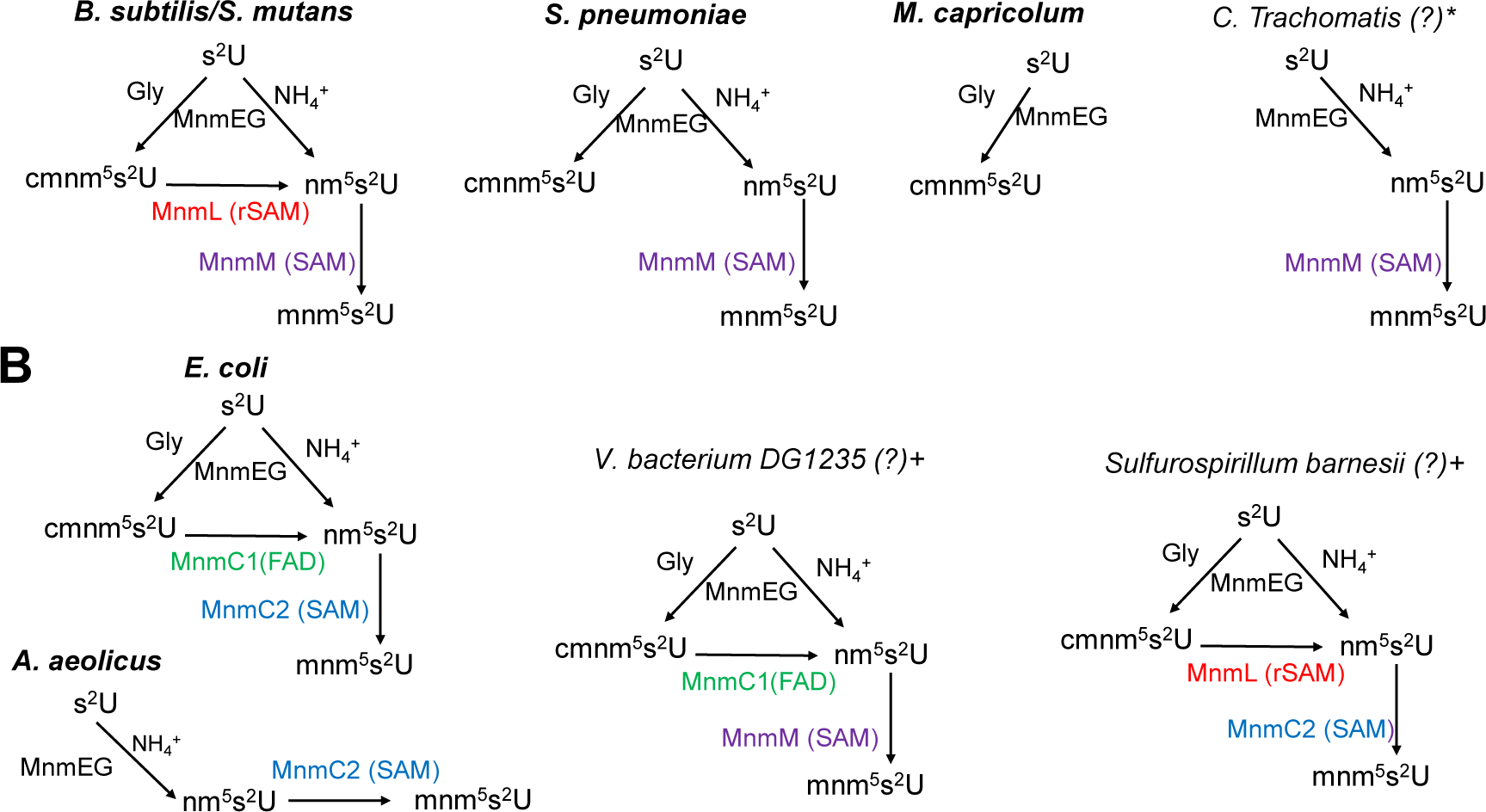
Diversity in the cnmn^5^s^2^U/mnm^5^s^2^U biosynthetic routes in bacteria. **(A)** Pathways utilizing MnmL(YtqA) and MnmM show varying complexity. **(B)** Bacteria utilize alternate enzymes or domains to complete the synthesis of nmn^5^s^2^U. The pathways in organisms in bold have been experimentally validated (see main text for references), and those marked with * and + have been predicted based on phylogenetic distribution and gene fusion data, respectively.

The occurrence of alternate pathways for the synthesis of mnm^5^s^2^U cannot be simply explained by the existence of an optimized single enzyme catalyzing two consecutive reactions as in the case of MnmC or by the recruitment of a proposed MnmL-MnmM complex. In fact, nature has found multiple combinations to promote hypermodification of U34 tRNA involving both C2 and C5 positions, which guarantees fidelity in translation. Distinct pathways have been ascribed for s^2^U in prokaryotes, as well as distinct cytosolic and mitochondrial eukaryotic systems (21, 42, 43). As discussed in the introduction, for modifications in the C5 position, some species do not synthesize mnm^5^s^2^U and only accumulate cmnm^5^s^2^U modification with apparent no metabolic penalties. At the same time, other species have bypassed the cmnm^5^s^2^U intermediate by utilizing only the ammonia route through the direct formation of nm^5^s^2^U intermediate and an MnmC2 enzyme to complete the pathway, as in the case of *A. aeolicus* (9). Phylogenetic distribution and fusion analyses (Fig. 2 and 3 and Fig. S1 and Fig. S8 of Cho et al., (22)) revealed the patchwork of enzymatic solutions that have evolved in bacteria for the synthesis of the x^5^s^2^U34 modifications (summarized Fig. 8). Some of these solutions have been experimentally validated (in bold in Fig. 8), while others are predicted from the fusions analysis (annotated with a + sign in Fig. 8) or from phylogenetic distribution data (annotated with a * sign in Fig. 8). For example, *Chlamydiae trachomatis* encodes an MnmM but no MnmL(YtqA) nor MnmC1 (Table S6 of Cho et al. (22) and Supplemental data S4B). We predict that, like *A. aoelicus*, this organism only uses the ammonium route to produce nm^5^s^2^U but then uses MnmM (instead of MnmC2 in *A. aoelicus*) to perform the last step (Fig. 8). Similarly, the MnmL/MnmC2 and MnmC1/MnmM fusions (Supplemental data S4A) suggest these two combinations of enzymes can be used to convert cmn^5^s^2^U into nm^5^s^2^U in addition to the known *E. coli* MnmC1/nmC2 combination. These predictions await experimental validation as one cannot rule out that other yet-unidentified enzymes are present in some organisms. Also, in organisms that only harbor MnmAGE, it is impossible to predict without experimental data if the glycine route or the NH_4_ route are used alone or in combination, even if to date no organisms seem to stop at the nm^5^s^2^U intermediate. It is also clear that the genes encoding enzymes involved in the late steps of the mnm^5^s^2^U synthesis pathway can be quite easily lost in specific clades, as shown here in the *S. pneumoniae* clade.

Functional assignment solely based on sequence similarities is not a suitable approach for computational prediction of the mnm^5^s^2^U pathways. Both YtqA/MnmL and YtqB/MnmM proteins are members of protein superfamilies that have many paralog subgroups and one must be very cautious in transferring functional annotations from the experimentally validated members to homologs in order to avoid mis-annotation of paralogs. The experimentally validated *B. subtills* MnmM is a member of the S-adenosyl-L-methionine-dependent methyltransferase superfamily (IPR029063) that contains hundreds of families and cannot be used to propagate the annotation. A more specific profile has been recently created, which places MnmM in the methyltransferase MnmM-like family (IPR010719), and it has been used to propagate the annotation in Interpro and Uniprot databases and experimentally validates for several members of the family (22). For the transfer of the MnmL/YtqA functional annotation, one can confidently propagate within the proteins in Group 1 of the SSN generated in Fig. 4. Proteins of subgroups 3, 7, and 8 also have a high probability of isofunctionality with proteins of subgroups 1 based on fusion and physical clustering (Fig. 4). For all other subgroups experimental testing is required before any annotation transfer can occur. One subgroup that is most certainly not isofunctional with MnmL/YtqA is group 4, which contains the *E. coli* YhcC protein. As discussed above, there is no cmnm^5^s^2^U to mm^5^s^2^U transformation activity when the *mnmC* gene of *E. coli* that encodes the mnmC1/MnmC2 fusion is deleted. In addition, physical clustering data of the larger YtqA/YhcC family never links members of *E. coli* YhcC subgroups with any protein of the nm^5^s^2^U pathway (Supplementary data S5). The genes that cluster most frequently with genes encoding proteins of the YhcC subgroup (group 4 in Fig. 4) are genes involved in anaerobic/aerobic growth conditions and oxidative stress resistance such as *gltB*, *gtlD*, *arcB*, *ptsN* and *elbB* (Supplementary data S5). It has also been shown that *yhcC* expression is regulated by the anaerobic transition regulator FNR (44).

Overall, this study shows the diversity of pathways involving the synthesis of xnm^5^s^2^U tRNA in bacteria. Genomic analysis guided the computational assignment of alternate pathways and led to the identification of MnmL/YtqA as a component of this pathway. The occurrence of fusion proteins containing alternate arrangements of MnmC1/MnmL and MnmC2/MnmM supports the proposed role of MnmL in catalyzing the conversion of cmnm^5^s^2^U into nm^5^U in a reaction that is coupled to the final methylation by MnmM. Experimental validation of such computational assignments were completed for both *S. mutans* and *B. subtilis*. Interestingly, analysis of *S. pneumoniae* modified tRNA suggests that the arrangement of MnmL as two polypeptides spanning the N- and C-halves of the enzyme leads to loss of function. Thus, *S. pneu*m*oniae* likely represents an evolutionary intermediate in route to losing this branch of the pathway. The accumulation of cmnm^5^s^2^U appears to compensate for the deficiency of the fully modified mnm^5^s^2^U, as strains lacking this modification do not display a growth phenotype or any other metabolic defects. Validation of functional assignments for alternate biosynthetic pathways awaits experimental demonstration. Nevertheless, the examples reported in this study provide direction for these investigations.

## Methods

### General bioinformatic resources

For literature and sequence retrievals, the resources at NCBI (https://www.ncbi.nlm.nih.gov/) (45), UniProt (https://www.uniprot.org) (46), and BV-BRC (https://www.bv-brc.org) (47) were routinely used. PaperBlast was used to find published papers on members of a given family (papers.genomics.lbl.gov/) (48). Sequence Similarity Networks (SSNs) were generated with the Enzyme Function Initiative (EFI) suite of webtools (https://efi.igb.illinois.edu/) (49). SSNs were visualized using Cytoscape (50). Species phylogenetic trees were usually constructed using phyloT v2 (data version 2023.2). Both species and protein phylogenetic trees were visualized and annotated using iTOL v6 (version 6.8.1; https://itol.embl.de) (51). The tree building and visualization tools of BV-BRC were also used. Absence-presence data were retrieved from the Database of Clustered Orthologous Genes (COG Db) (https://ftp.ncbi.nih.gov/pub/COG/COG2020/data/, accessed 22 August 2022) (34). Physical clustering data were retrieved from the KEGG Database (via protein sequence entry option, “ene luster”). Metabolic pathway and gene symbol/family name annotations were retrieved from KEGG (release 108.0; https://www.kegg.jp/kegg/) (52) and UniProtKB (release 2023_04; https://www.uniprot.org) (46). Gizmogene (http://www.gizmogene.com/) was used to create gene cluster figures. SubtiWiki was used to explore coexpression data in *B. subtilis* (53). Protein sequences were aligned using MultiAlin (54).

### Absence-presence data extraction, benchmark genome filtration for encoding of MnmAGE homologs, and MnmC1-/MnmC2fusion analysis

The MnmAGE families (COG0482, COG0445, and COG0486, respectively) absence-presence data was extracted from the Database of Clustered Orthologous Genes (COG) (https://ftp.ncbi.nih.gov/pub/COG/COG2020/data/, accessed 22 August 2022) (34), This set was used to create a benchmark set of 968 bacterial genomes encoding MnmAGE homologs (Supplemental data S1) that was used to then analyze the presence/absence of MnmC protein family members (COG4121). Because COG does not distinguish between the different subgroups of MnmC (i.e., MnmC1, MnmC2, MnmC1/C2 fusion), CDD Search (batch search, standard view) was used to determine MnmC1 domain-containing proteins, MnmC2 domain-containing proteins, MnmC1/C2 fusions, and other fusions with MnmC2 (https://www.ncbi.nlm.nih.gov/Structure/bwrpsb/bwrpsb.cgi, accessed 22 August 2022) (55). The specific hits per CDD Search results provided the labels of sequences used in data tables and figures (Supplemental data S3). Batch CDD search results also supplied respective sequence lengths.

### Physical clustering analyses

Using the COG database for the 968 bacterial benchmark genomes filtered for MnmAGE, YtqA family sequences were acquired (ordered locus names per assembly) (Supplemental Data S4). For each identified YtqA homolog, the KEGG Database “ ene luster” protein entry feature was used to extract physical clustering data. ach set of genes per “ ene luster” varied in number, determined by KEGG. These data were expanded into a tabular format per respective cluster YtqA sequence (UniProt Accession ID) (Supplemental data S5). All gene cluster proteins were mapped to their respective UniProt Accession IDs and investigated for methyltransferase (i.e., MnmM-/YtqB-like) domains using three different methods, one BLAST-driven and two HMM-driven (Supplemental data S5C-E): 1) traditional BLASTp (release, ncbi-blast-2.14.1+-win64; e-value = 0.0001; UniProtKB derived FASTA sequences as inputs) (56); 2) CDD sequence search (output format, Standard; RefSeq identifiers as inputs) (57); and 3) InterProScan sequence search (UniProtKB derived FASTA sequences as inputs) (58). All MnmM-/YtqB-like absence-presence data was managed, curated, and the outputs of the three different retrieval methods were merged using Microsoft Office Excel (Supplemental data S5C and 5D). The same was true for the YtqA-containing gene cluster data and subsequent data transformations required for calculating absence-presence proportions/percentages across various genome subgroups within phylum-level defined taxonomic groups (taxonomic groups derived from those included in the COG absence-presence data; representative taxonomic group tax IDs were manually curated) (Supplemental data S6). These mapped percentages would be the source data mapped to the phylogenetic trees of Figure 2. Trees were generated, mapped, and edited using phyloT v2 (https://phylot.biobyte.de/index.cgi) and iToL (51). UniProt IDs for all genes of each cluster were mapped. Metabolic categories were assigned to each cluster member using KEGG and UniProt pathway category assignments (Supplemental data S8). Gene symbol/family names were assigned per their respective annotations (Supplemental data S8). These data, annotations were subsequently mapped to tree using iTOL or to SSNs using Cytoscape.

### Sequence similarity and gene neighborhood networks

Sequence Similarity Networks (SSNs) were generated with the Enzyme Function Initiative (EFI) suite of webtools (https://efi.igb.illinois.edu/) (49). SSNs were visualized using Cytoscape (50). A network was generated using the InterPro identifier IPR005911 (Protein YhcC-like), taxonomy restricted to include only bacterial sequences. Sequences annotated as fragments by UniProt were excluded. Three preliminary networks were generated for alignment scores (AS) of 90, 97, and finally 100 (‘repnode’ percentages of 95, 95, and 55, respectively). AS100, repnode 55 was selected for figure generation. After acquiring UniProt accession IDs for all sequences possible, fusions and absence-presence data for MnmC and YtqA (see preceding subsection) were mapped to each preliminary network and then to the final colorized network (Fig. 4). The AS100 network was colorized by cluster and with all singletons and 2-node clusters removed (Supplemental data S9).

### Density functional theory (geometry optimizations)

Density functional theory (DFT) calculations were carried out using Orca 5.0 (59)The set of cmnm^5^s^2^U molecular models was generated with a variety of protonation states and sites for the initial H-atom abstraction (summarized in Fig.S6) using Chem3D (PerkinElmer). The ribose moiety and the rest of the tRNA were approximated with a methyl group, but all other atoms of the 2-thiouracil were kept as shown in Fig. S6A. Geometry optimization was done in Orca with the BP86 functional (60, 61) and def2-SVP basis set (62) with def2/J auxiliary functions (63) and using the conductor-like polarizable continuum model for water (dielectric constant = 80.4, refractive index = 1.33)(64).

### Bacterial Strains and Growth Conditions

All bacterial strains and plasmids used in this work are listed in Table S3. Strains of *S. mutans* were routinely cultured statically at 37°C in 5% CO_2_ in BHI medium (BD Biosciences) with 10 µg/mL erythromycin (Sigma) when appropriate. Strains of *B. subtilis* were routinely cultured with shaking at 200 rpm in LB media (Fisher) at 37 °C with 50 mg/mL of kanamycin (Sigma-Aldrich) or 10 µg/mL erythromycin when appropriate. *E. coli* strains were cultured with shaking at 200 rpm on LB agar at 37°C supplemented with 100 µg/mL ampicillin (Acros) or 50 µg/mL kanamycin when appropriate. *S. pneumoniae* cells were cultured statically in BHI medium at 37°C in an atmosphere of 5% CO_2_. *S. pneumoniae* were strains grown on plates containing trypticase soy agar II (modified; Becton-Dickinson), and 5% (vol/vol) defibrinated sheep blood (TSAII-BA). Plates were incubated at 37°C in an atmosphere of 5% CO_2_. TSAII-BA plates for selections contained antibiotics at concentrations described previously (65).

### Genetic constructions methods

#### E. coli

The *B. subtilis ytqA-mnmM* gene region was amplified from *B. subtilis* CU1065 genomic DNA prepared using a DNA extraction kit (QuickExtract, Epicentre). PCR amplification was performed using primers YtqA_F and YtqB_R (Table S3) and the resulting fragment was digested with NcoI and BamHI prior to ligation into NcoI/BglII sites of pBAD, the resulting plasmid was named pDS358. The coding sequence for *ytqA* and *ytqB* were amplified with restriction sites NcoI and XhoI/BamHI and NcoI and amHI from pDS358 by using primers YtqA_NcoI_F and YtqA_XhoIBamHI_R and YtqB_NcoI_F and YtqB_BamHI_R respectively. The resulting amplicons were digested and ligated into pBAD to create pDB359 and pDS365, respectively. The sequences of these constructs were verified through DNA sequence (Azenta). pBAD expression plasmids containing *ytqA-mnmM*, *ytqA*, *mnmM* were transformed into *E. coli* Δ*mnmC* and cultured in 500 m of B medium containing ampicillin (100 μg/m) and arabinose (2 mg/m) up to OD600 nm of ∼1.0. Cells were harvested by centrifugation and stored at -20oC prior to further analysis.

#### S. mutans

Mutations in *S. mutans* UA159 were constructed by a PCR ligation mutagenesis method (66)). Different sets of primers (listed in Table S4) were used to amplify regions directly upstream and downstream from the gene targeted for deletion and to introduce complementary ligation sites to a non-polar erythromycin resistance cassette. The upstream, downstream, and resistance markers were digested and ligated into a linear piece of DNA, which was subsequently used to transform *S. mutans* cells in the presence of competence stimulating peptide (67) and double-cross-over recombination events were selected by plating on BHI agar supplemented with 10 µg/mL erythromycin. The correct deletion of the gene was confirmed by PCR with external primers and amplicon length comparison.

#### S. pneumoniae

*Streptococcus pneumoniae* strains used in this study are listed in Table S3. Strains were derived from unencapsulated strain IU1824 (D39 Δ*cps rpsL1*), which were derived from the encapsulated serotype-2 D39W progenitor strain IU1690 (68). Compared with the progenitor strain IU1690, a spontaneous mutation (Q150STOP) was found in *spd_0567* (*mnmM*) during the construction of IU1824 (69). The *mnmM* (Q150STOP) mutation in IU1824 was repaired to *mnmM*^+^ in strain IU19835. No growth or microscopic differences were observed between IU1824 and IU19835 (*spd_0567*^+^) in BHI broth. Strains containing antibiotic markers were constructed by transformation of CSP1-induced competent pneumococcal cells with linear DNA amplicons synthesized by overlapping fusion PCR (65). Strains containing markerless alleles in native chromosomal loci were constructed using allele replacement via the P_c_-[*kan*-*rpsL*^+^] (70). Primers used to synthesize different amplicons are listed in Table S4. TT1519, TT1522 are outside forward primers and TT1520 is the outside reverse primer for strains IU19964, IU19966 and IU19968. Fusion primers for IU19964, IU19966 and IU19968 are TT1618/TT1619, TT1623/TT1624 and TT1621/TT1622, respectively. D39 genomic DNA was used as a template. For the construction of IU19974, 5’ fragment was obtained using IU11077 (66) which contains 5’ fragment of Δ*bgaA*::*kan*-t1t2-P_Zn_ and primers TT657 and TT1577. Middle fragment and 3’ fragments were obtained using D39 gDNA as templates and primer pairs TT1578/TT1579 and TT1580/CS121. Mutant constructs were confirmed by PCR and DNA sequencing of chromosomal regions corresponding to the amplicon regions used for transformation. Ectopic expression of *mnmM* was achieved with a P_Zn_ zinc-inducible promoter in the ectopic *bgaA* site. 0.2 mM Zn^2+^ and 0.02 mM Mn^2+^ were added to BHI broth for inducing conditions. Mn^2+^ was added with Zn^2+^ to prevent zinc toxicity (71).

#### B. subtilis

The *B. subtilis mnmM* and *ytqA* mutant were obtained from the Bacillus Genetic Stock Center (https://bgsc.org/index.php) and verified by PCR using Oligos BSU_ytqB_VF and BSU_ytqB_VR or BSU_ytqA_VF and BSU_ytqA_VR (see Table S2).

### Bulk tRNA Extraction

Samples of bacterial culture were used to prepare bulk tRNA extracts for *E. coli*, *S. mutans*, *S. pneumoniae*, and *B. subtilis.* As the LC/MS analyses were done by three laboratories and utilized different methods to prepare tRNA samples were used. In the PDS method, 0.5 L cultures of *E. coli* and *B. subtilis* were harvested by centrifugation, and the cell pellets were treated as previously described to obtain the bulk tRNA samples (72) that were stored at -20°C until LC-MS analysis. In the VDCL1 method*, B. subtilis* cell pellets were prepared by diluting overnight (16-18 hours) liquid cultures to an initial OD_600nm_ of 0.02, allowed to grow at 37 °C with shaking at 200 rpm until a final OD_600nm_ of ∼1.0 or another specified value as indicated, and 5 ml of cells were centrifuged. For *B. subtilis* samples, each cell pellet was re-suspended in a solution containing 10 mg/mL lysozyme and sodium acetate (pH5.3, 50 mM sodium acetate) and transferred to a 1.5 mL microcentrifuge tube followed by an incubation at 37°C for 1 hour. The resulting lysate was then used to isolate bulk tRNA as previously described (73).

*S. mutans* tRNA samples were prepared using the VDCL2 method. Using this method, each sample was prepared by first diluting overnight (16-18 hours) liquid cultures to an initial OD_600nm_ of 0.02 and allowed to grow statically at 37°C in 5% CO_2_ until a final OD_600nm_ of ∼0.6 was achieved, and 5 ml were centrifuged. Each *S. mutans* cell pellet was re-suspended in a solution 5 mg/mL lysozyme (lysozyme from chicken egg white, Sigma Aldrich - L6876) and TE Buffer (pH8, 10 mM Tris, 1mM EDTA) and transferred to a microcentrifuge tube. Each sample was then incubated on wet ice for 30 minutes. After the cold incubation, a solution of 10% SDS in water was added to each sample tube to reach a final concentration of 1% SDS, followed by an incubation at 95°C for 5 minutes. The resulting *S. mutans* lysate was then used to isolate bulk tRNA as previously described (73).

*S. pneumoniae* tRNA samples were prepared using the MW method. Each sample processed by the MW was prepared by diluting overnight cultures (12-15 hours) to an initial OD_620nm_ 0.005 and allowed to grow until an OD_620nm_ of 0.1 - 0.3 was achieved. For induction of *mnmM* in the merodiploid strain IU19974, 0.2 mM Zn^2+,^ and 0.02 mM Mn^2+^ were added to BHI broth during day growth. For each tRNA preparation, two 40 mL of culture grown to OD_620_ ≈0.19 to 0.22 were centrifuged at 16,000 x *g* for 7 min at 4°C. Each pellet was added with 100 µL of lytic buffer (1.2 % triton, 2 mM EDTA, 20 mM Tris, pH 8.3), vortexed, and followed by the addition of 500 µL of Trizol. The bacteria Trizol mixtures from the two pellets were combined and added to a 2-mL tube containing lysing matrix B (MP biomedical, 116911-500), and processed with a FastPrep-24 machine (MP biomedical) for three times for 40 sec at setting 6 with a 24×2 holder at 4°C. The tubes were centrifuged at 16,000 x *g* for 7 min at 4°C, and the upper layers (∼900 µL) were transferred to new microfuge tubes and incubated for 5 min at room temperature. 450 µL of chloroform was added to the transferred sample, vortexed for 10 sec and incubated for 5 min at room temperature, followed by centrifugation at 16,000 x *g* for 10 min at 4°C. Approximately 450 µL of the upper phase was transferred to microfuge tubes, added with 0.5 x volume (225 µL) of 100 % room temperature ethanol, and inverted 10 times to mix. The mixture was added to a PureLink miRNA Isolation Kit Spin Columns, followed by centrifugation at 1 min at 12,000 x *g*. The flow through (∼600 µL) was transferred to a new microfuge tube and mixed with 700 µL of 100 % ethanol. The mixture was added in 2 runs to a new PureLink miRNA Isolation Kit Spin Column, centrifuged for 1 min at 12,000 g, and the flow-through discarded. Wash buffer W5 from the PureLink miRNA Isolation Kit (500 µL) was added to the column and centrifuged for 1 min at 12,000 x *g.* The wash step was performed 3 times, followed by a final centrifugation of 3 min. The spin column was transferred to a new microfuge tube, and 62 µL of RNase-free water was added and incubated for 1 min. The sample was eluted by centrifugation for 1 min at 21,000 x *g*. Samples were quantified with a NanoDrop spectrophotometer, showing typical 260/280 readings of ∼2 and 260/230 readings of 2.0 to 2.2. Typical yields from this protocol were 40 to 70 µg from 80 mL of cell culture collected at OD_620_ ≈0.19 to 0.22. The purified samples were stored at -80°C until further analysis.

### Nucleoside preparation and mass spectrometry analysis for *B. subtilis*, *E. coli*, *S. pneumoniae* tRNA samples (LC-MS Method 1)

Forty micrograms of total RNA were heat-denatured at 100 °C for 5 minutes, chilled on ice for 5 minutes, and allowed to come to room temperature. The RNA was digested in 20 mM TMA-acetate pH 5.3 containing 5 µL P1 nuclease, 10 µL alkaline phosphatase (RSAP) and 20 µL CutSmart buffer. The reaction was incubated at 37°C for 1 hour, centrifuged at 12,800 *g* for 10 minutes, and the supernatant was spiked with 5 µL of methanol-formic acid (80:4) before mass spectrometry analysis. Digested total RNA samples were analyzed by HPLC using a Polaris C-18 column (Agilent), coupled with a mass spectrometer (Orbitrap) in positive ion mode, as described in Edwards et al. (74). Optima LCMS grade solvents were 0.1% formic acid in water (solvent A) and 0.1% formic acid in methanol (solvent B). The column was pre-equilibrated with 2% Solvent B for 15 column volumes at 300 μ min^-1^. Tune file parameters used during runs were as follows: Voltage (kV): 4.01; Sheath Gas Flow Rate (arb): 47.00; Auxiliary Gas Flow Rate (arb): 30.00; Sweep Gas Flow Rate (arb): 0.00; Capillary Voltage (V): 2.00; Capillary Temperature (C): 350.00; Tube Lens Voltage (V): 49.89. Digested nucleosides were separated and detected over the course of a 39-min run using the gradient of solvent B as follows: 0–4 min, hold at 2% B (v/v); 4–25 min, increase B from 2% to 100%; 25–33 min, hold at 100% B; 33–39 min switch from 100% B to 2% B to allow column re-equilibration at 2% B. Standards of dihydrouridine (Dalton) and pseudouridine (Barry & Associates) were run as controls for normalizing data. The desired analytes D ([M+H]^+^:247.0925), nm^5^s^2^U ([M+H]^+^: 290.0805), cmnm^5^s^2^U ([M+H]^+^: 348.086), mnm^5^s^2^U ([M+H]^+^:304.096), where the masses were determined within 5 ppm accuracy.

### Nucleoside preparation and mass spectrometry analysis of *B. subtilis* and *S. mutans* samples (LC-MS Method 2)

tRNA from each sample (*B. subtilis:* 1 µg; *S. mutans* 2 µg) was hydrolyzed in a 50 µL digestion cocktail containing 12.5 U (*B. subtilis*) or 8 U (*S. mutans*) benzonase, 5 U CIAP (calf intestinal alkaline phosphatase), 0.15 U PDE I (phosphodiesterase I), 0.1 mM deferoxamine, 0.1 mM BHT (butylated hydroxytoluene), 5 ng coformycin, 50 nM 15N-dA (internal standard [^15^N]_5_-deoxyadenosine), 2.5 mM MgCl_2_ and 5 mM Tris-Hcl buffer pH 8.0. The digestion mixture was incubated at 37 °C for 6 h. After digestion, all samples were analyzed by chromatography-coupled triple-quadrupole mass spectrometry (LC-MS/MS). For each sample, 200 ng (*B. subtilis*) or 600 ng (*S. mutans*) of hydrolysate was injected in two technical replicates. Using synthetic standards, HPLC retention times of RNA modifications were confirmed on a Waters Acuity BEH C18 column (50 × 2.1 mm inner diameter, 1.7 µm particle size) coupled to an Agilent 1290 HPLC system and an Agilent 6495 triple-quad mass spectrometer. The Agilent sample vial insert was used. The HPLC system was operated at 25 °C and a flow rate of 0.3 mL/min in a gradient (Table S5) with Buffer A (0.02% formic acid in double distilled water) and Buffer B (0.02% formic acid in 70% acetonitrile). The HPLC column was coupled to an Agilent 6495 triple quad mass spectrometer with an electrospray ionization source in positive mode with the following parameters: Dry gas temperature, 200 °C; gas flow, 11 L/min; nebulizer, 20 psi; sheath gas temperature, 300 °C; sheath gas flow, 12 L/min; capillary voltage, 3000 V; nozzle voltage, 0 V. Multiple reaction monitoring (MRM) mode was used for detection of product ions derived from the precursor ions for all the RNA modifications with instrument parameters including the collision energy (CE) optimized for maximal sensitivity for the modification. Based on synthetic standards (Biosynth) with optimized collision energies, the following transitions and retention times were monitored: cmnm^5^s^2^U, m/z 348 → 141, 2.3 min; mnm^5^s^2^U, m/z 304 → 172, 1.6 min; nm^5^s^2^U, m/z 290 → 158, 1.2 min; D, m/z 115, 0.864 min. Signal intensities for each ribonucleoside were normalized by dividing by the sum of the UV signal intensities of the four canonical ribonucleosides recorded with an in-line UV spectrophotometer at 260 nm.

## Acknowledgments.

This work was funded by the National Institutes of Health (R01 GM70641 to V. dC-L. and R35 GM131767 to MEW and ES026856, and ES024615 and ES031529 to Thomas Begley), the National Research Foundation of Singapore through the Singapore-MIT Alliance for Research and Technology Antimicrobial Resistance Interdisciplinary Research Group (PCD) and the and National Science Foundation (MCB-1716535 to PSD). We would like to thank Chelsey Bomar for identifying the first YtqA_MnmD fusion during the MCB6318 Spring 2022 class and Scott Dawson, Brystol Habermacher, and Mira Baki for assistance in the cloning of *B. subtilis* genes.

