## Supplemental_Tables_and)Figures for "Alternate routes to mnm^5^s^2^U synthesis in Gram-positive bacteria"

**Table S1.** Distribution of MnmC2, MnmC1/C2 fusions, YtqA, and instances of YtqA homologs physically clustered with YtqB.

| | | | | | | | **Of Those w/o MnmC** (MnmC1, MnmC2, or MnmC1/C2 fusion) | |
|---|---|---|---|---|---|---|---|---|
| **Group Name** | **Tax ID** | **Number of Genomes** | **YtqA (%)** | **YtqA clusters w. YtqB (%)** | **MnmC2 (%)** | **MnmC1/ C2 Fusion (%)** | **YtqA (%)** | **YtqA clusters w. YtqB (%)** |
| **Acidobacteria** | 57723 | 7 | 0 | NA | 0 | 0 | 0 | 0 |
| **Aquificae** | 187857 | 9 | 89 | 13 | 78 | 0 | 100 | 50 |
| **Bacteroidetes** | 200643 | 106 | 23 | 13 | 85 | 0 | 25 | 6 |
| **Chlamydiae** | 204428 | 6 | 0 | NA | 0 | 0 | 0 | 0 |
| **Chlorobi** | 1090 | 5 | 0 | NA | 0 | 0 | 0 | 0 |
| **Chloroflexi** | 32061 | 5 | 0 | NA | 0 | 0 | 0 | 0 |
| **Cyanobacteria** | 1117 | 41 | 5 | 0 | 90 | 0 | 25 | 0 |
| **Deferribacteres** | 68337 | 5 | 100 | 40 | 0 | 0 | 0 | 0 |
| **Deinococcus-thermus** | 1297 | 6 | 0 | NA | 0 | 0 | 0 | 0 |
| **Bacilli** | 91061 | 67 | 45 | 80 | 0 | 0 | 67 | 36 |
| **Clostridia** | 186801 | 75 | 60 | 22 | 0 | 0 | 71 | 13 |
| **Negativicutes** | 909932 | 10 | 60 | 83 | 0 | 0 | 70 | 50 |
| **Tissierellia** | 1737404 | 8 | 75 | 17 | 0 | 0 | 88 | 13 |
| **Other Firmicutes** | 53420* | 3 | 33 | 100 | 0 | 0 | 67 | 33 |
| **Fusobacteria** | 203490 | 6 | 50 | 0 | 0 | 0 | 50 | 0 |
| **Mollicutes** | 31969 | 14 | 14 | 50 | 0 | 0 | 14 | 7 |
| **Planctomycetes** | 112 | 14 | 29 | 0 | 0 | 0 | 29 | 0 |
| **Alphaproteobacteria** | 28211 | 150 | 0 | NA | 31 | 10 | 0 | 0 |
| **Betaproteobacteria** | 28216 | 99 | 12 | 33 | 4 | 65 | 3 | 0 |
| **Deltaproteobacteria** | 28221 | 32 | 47 | 47 | 19 | 0 | 69 | 27 |
| **Epsilonproteobacteria** | 3031852 | 12 | 67 | 25 | 75 | 8 | 3 | 0 |
| **Gammaproteobacteria** | 1236 | 222 | 18 | 41 | 3 | 60 | 1 | 0 |
| **Other Proteobacteria** | 3018035 | 6 | 17 | 100 | 50 | 17 | 0 | 0 |
| **Spirochaetes** | 203692 | 10 | 40 | 0 | 50 | 10 | 75 | 0 |
| **Synergistetes** | 508458 | 5 | 40 | 0 | 0 | 0 | 40 | 0 |
| **Thermotogae** | 188708 | 9 | 56 | 40 | 0 | 0 | 78 | 22 |
| **Verrucomicrobia** | 74201 | 9 | 11 | 0 | 33 | 0 | 17 | 0 |
| **Other Bacteria** | 2* | 27 | 30 | 25 | 15 | 0 | 26 | 9 |

**Taxonomic clades not included in iToL trees*

**Table S2.** DFT-computed geometric values[a] for Models of cmnm$^5$s$^2$U following H-atom abstraction

| Model | $C_c$—$C_d$ (Å) | $N_b$—$C_c$ (Å) | $C_a$—$N_b$ (Å) |
|---|---|---|---|
| **UHH**[b] | 1.533 | 1.487 | 1.508 |
| $C_c^\bullet$ | 1.459 | 1.422 | 1.516 |
| $N_b^\bullet$ | 1.524 | 1.431 | 1.428 |
| $C_a^\bullet$ | 1.524 | 1.501 | 1.458 |
| **UH** | 1.546 | 1.470 | 1.467 |
| $C_c^\bullet$ | 1.444 | 1.371 | 1.454 |
| $N_b^\bullet$ | 1.855 | 1.372 | 1.450 |
| $C_a^\bullet$ | 1.540 | 1.454 | 1.374 |
| **U** | 1.592 | 1.461 | 1.511 |
| $C_c^\bullet$ | 1.486 | 1.384 | 1.445 |
| $N_b^\bullet$ | 1.677 | 1.401 | 1.435 |
| $C_a^\bullet$ | 1.570 | 1.449 | 1.345 |

a. Bond lengths shown in blue are seen to shorten significantly upon the denoted H-atom abstraction, indicating partial double bond formation. Bond lengths shown in red are seen to lengthen indicating bond breaking.

b. Geometric parameters listed for models named **UHH**, **UH**, and **U** are those for cmnm$^5$s$^2$U models *prior* to H-atom abstraction.

**Table S3. Strains and plasmids used.**

| Strain, plasmid | Phenotype, genotype and/or description | Ref/Source |
|---|---|---|
| *S. pneumoniae* | | |
| IU1824 | D39 rpsL1 Δ*cps rpsL1 spd_0567*(*mnmM*)(Q152stop) | (1) |
| IU19821 | D39 Δ*cps rpsL1* Δ*spd_0567*::$P_c$-[*sacB-kan-rpsL*$^+$] | (2) |
| IU19835 | D39 Δ*cps rpsL1 spd_0567*$^+$ | (2) |
| IU19964 | D39 Δ*cps rpsL1* Δ[*spd_0565-0566*] (Δ*ytqA*) markerless (IU19821 X fusion Δ[*spd_0565-0566*] amplicon) | This study |
| IU19966 | D39 Δ*cps rpsL1* Δ*spd_0567* (Δ*mnmM*) markerless (IU19821 X fusion Δ*spd_0567* amplicon) | This study |
| IU19968 | D39 Δ*cps rpsL1* D39 Δ*cps rpsL1* Δ[*spd_0565-0567*] (Δ*ytqA mnmM*) markerless (IU19821 X fusion Δ[*spd_0565-0567*] amplicon) | This study |
| IU19974 | D39 Δ*cps rpsL1* Δspd_0567 (Δ*mnmM* markerless//Δ*bgaA*::*kan*-t1t2-$P_{Zn}$-*mnmM*$^+$) (IU19966 X fusion Δ*bgaA*::*kan*-t1t2-$P_{Zn}$-*mnmM*$^+$) | This study |
| *B. subtilis* | | |
| CU1065 | Wild-Type, *trpC2* attSPβ sfp0 | Laboratory Stock |
| BKE30480 | CU1056 Δ*ytqA*:: $Erm^R$ | (3) |
| BKE30490 | CU1056 Δ*mnmM*:: $Erm^R$ | (3) |
| *S. mutans* | | |

| | | |
|---|---|---|
| S. mutans UA159 | Serotype c, ATCC 700610 | (4) |
| MSJ1093 | *S. mutans* UA159 Δ*ytqA*::Erm$^R$ | This study |
| MSJ1095 | *S. mutans* Δ*ymnmM*::Erm$^R$ | This study |
| MSJ1097 | *S. mutans* Δ*ytqA mnmM*::Erm$^R$ | This study |
| *E. coli* | | |
| JW5380-2 | Δ(araD-araB)567, ΔlacZ4787(::rrnB-3), $\lambda^-$, ΔtrmC732::kan, rph-1, Δ(rhaD-rhaB)568, hsdR514 | (5) |
| TOP10 | F- mcrA Δ(mrr-hsdRMS-mcrBC) φ80 lacZΔM15Δ lacX74 recA1 araΔ139Δ(ara-leu)7697 galU galK rpsL (StrR) endA1 nupG | Invitrogen |
| **Plasmids** | | |
| pBAD pBAD/Myc-His | Amp$^R$, ColE1 arabinose inducible promoter | (6) |
| pDS358 | pBAD-*ytqAB*$_{Bsu}$ | This study |
| pDS359 | pBAD-*ytqA*$_{Bsu}$ | This study |
| pDS365 | pBAD-*ytqB*$_{Bsu}$ | This study |

**Table S4**. Oligonucleotides used

| Oligo | Sequence 5′ - 3′ |
|---|---|
| For construction of MSJ1093 | |
| ytqA P1 | GCAGTGATTGCTTTAGTTGTCC |
| ytqA P2 | GCGCGCTGCAGACAAAAAGTACAGCCACCATGA |
| ytqA P3 | GCGCGGAGCTCTCATTCATCGTCTGACAGGTGA |
| ytqA P4 | ATCCTCCTCCTTCGTAGAGGTA |
| For construction of MSJ1095 | |
| ytqB P1 | GTGATGGAACTGTGGCTCATG |
| ytqB P2 | GCGCGCTGCAGGGCCTGTTCCTGTACATCAAAA |
| ytqB P3 | GCGCGGAGCTCGGGTGGCGATAAGGAAAAAGAT |
| ytqBP4 | AGTCAAAAAGGGAGAAAATTGGGT |
| For construction of MSJ1097 | |
| ytqAB P1 | GCAGTGATTGCTTTAGTTGTCC |
| ytqAB P2 | GCGCGCTGCAGCGTACAAAAAGTACAGCCACCA |
| ytqAB P3 | GCGCGGAGCTCGGGTGGCGATAAGGAAAAAGAT |
| ytqAB P4 | AGTCAAAAAGGGAGAAAATTGGGT |
| For verification of MSJ1062 and MSJ1063 | |
| BSU_ytqB_VF | CAAAGAACTGCTGAAAATGGCC |
| BSU_ytqB_VR | TGCTGATTGATAAACCCGTATGT |
| BSU_ytqA_VF | GCACGAAAAATGGAAGGACG |
| BSU_ytqA_VR | GCCGTGTTTTCTTAATTTATTGACG |
| For verification of MSJ1093, MSJ1095, and MSJ1097 | |
| SMU_ytqA_VF | CCAGTGTGTTAAACTGGAATCAC |
| SMU_ytqA_VR | AGTCGCATCAATGGCTATACTG |
| SMU_ytqB_VF | CATTCATCGTCTGACAGGTGA |

| | |
|---|---|
| SMU_ytqB_VR | ATCCTCCTCCTTCGTAGAGGTA |
| For construction of IU19964 | |
| TT1519 | GGCATGATAGAAGGCGTAGAAGTGATGTTCCTCC |
| TT1618 | TTACAGCCTTGCATCCTTGAACACTTCCTTCTCCAAAGAGTTTTCGATAATAATCATT |
| TT1619 | AATGATTATTATCGAAAACTCTTTGGAGAAGGAAGTGTTCAAGGATGCAAGGCTGTAA |
| TT1520 | CCAGAAGCATCATTCAAGAGTCCTTCGCCC |
| For construction of IU19966 | |
| TT1522 | GTCCCTATTGATGCGGAATTTGACTGTCCC |
| TT1623 | AATCATCACTAAAAACGGCGGGTTGTTGACTCCCATAGTCGCATCCACTACGACATCCTC |
| TT1624 | GAGGATGTCGTAGTGGATGCGACTATGGGAGTCAACAACCCGCCGTTTTTAGTGATGATT |
| TT1520 | CCAGAAGCATCATTCAAGAGTCCTTCGCCC |
| For construction of IU19968 | |
| TT1519 | GGCATGATAGAAGGCGTAGAAGTGATGTTCCTCC |
| TT1621 | ATCATCACTAAAAACGGCGGGTTGTTGACTTCTCCAAAGAGTTTTCGATAATAATCATT |
| TT1622 | AATGATTATTATCGAAAACTCTTTGGAGAAGTCAACAACCCGCCGTTTTTAGTGATGAT |
| TT1520 | CCAGAAGCATCATTCAAGAGTCCTTCGCCC |
| For construction of IU19974 | |
| TT657 | CGCCCCAAGTTCATCACCAATGACATCAAC |
| TT1577 | TGCCATCTCAAGTGGTCTTTTCATTTTCAATACATCGCTTCCTCTCTATCTTCCTTGTTA |
| TT1578 | TAACAAGGAAGATAGAGAGGAAGCGATGTATTGAAAATGAAAAGACCACTTGAGATGGCA |
| TT1579 | ACTGGTTTATGAGAAAGTAAGTTCTTTTATCCATGTCTGTATCTCTCTAATTTTTCAATC |
| TT1580 | AAAAATTAGAGAGATACAGACATGGATAAAAGAACTTACTTTCTCATAAACCAGTTGCTG |
| CS121 | GCTTTCTTGAGGCAATTCACTTGGTGC |
| For Construction of pBAD24 Complementation Vectors | |
| YtqA_NcoI_F | GTACCATGGAGAACAATCCTTTTCCTTATTCAAATAC |
| YtqA_XhoIBamHI_R | ACTGGATCCTCACTCGAGCGCTGATTCCTCCTCAAGCCG |
| YtqB_BamHI_R | GACTGGATCCTCATTTGCTGATCTGAGCTTTT |
| YtqB_NcoI_F | GTACCATGGTTTTGAAGAAAATTCTTCCTTACAGCAA |

**Table S5.** Buffer gradient used in the LC-MS/MS Method 2. Buffer A: 0.02% Formic acid in double distilled water. Buffer B: 0.02% Formic acid in 70% Acetonitrile.

| Time (min) | Buffer A (%) | Buffer B (%) | Flow (ml/min) |
|---|---|---|---|
| 0 | 100 | 0 | 0.3 |
| 5 | 99 | 1 | 0.3 |
| 6 | 98 | 2 | 0.3 |
| 7 | 97 | 3 | 0.3 |
| 8 | 95 | 5 | 0.3 |
| 9 | 93 | 7 | 0.3 |
| 10 | 90 | 10 | 0.3 |
| 12 | 88 | 12 | 0.3 |
| 13 | 85 | 15 | 0.3 |
| 15 | 80 | 20 | 0.3 |
| 16 | 25 | 75 | 0.3 |
| 17 | 0 | 100 | 0.3 |
| 18 | 0 | 100 | 0.3 |
| 20 | 0 | 100 | 0.3 |
| 21 | 100 | 0 | 0.3 |
| 25 | 100 | 0 | 0.3 |

**Table S6.** Identifiers for proteins and genomes used in Fig. S4 and S5

| YtqA_Name in FigS4A | Genome | Genome ID | Accession | BRC ID | RefSeq Locus Tag | Protein ID | AA length |
|---|---|---|---|---|---|---|---|
| N_Spneumoniae_ST556 | Streptococcus pneumoniae ST556 | 1130804.3 | CP003357 | fig\|1130804.3.peg.715 | MYY_0697 | AFC94358.1 | 169 |
| C_Spneumoniae_ST556 | Streptococcus pneumoniae ST556 | 1130804.3 | CP003357 | fig\|1130804.3.peg.716 | MYY_0698 | AFC94359.1 | 64 |
| N_Spneumoniae_TIGR4 | Streptococcus pneumoniae TIGR4 | 170187.11 | NC_003028 | fig\|170187.11.peg.680 | | | 169 |
| C_Spneumoniae_TIGR4 | Streptococcus pneumoniae TIGR4 | 170187.11 | NC_003028 | fig\|170187.11.peg.681 | | | 148 |
| N_Spneumoniae_R6 | Streptococcus pneumoniae R6 | 171101.6 | NC_003098 | fig\|171101.6.peg.635 | spr0569 | NP_358163.1 | 150 |

| | | | | | | | |
|---|---|---|---|---|---|---|---|
| C_Spneumoniae_R6 | Streptococcus pneumoniae R6 | 171101.6 | NC_003098 | fig\|171101.6.peg.636 | spr0570 | NP_358164.1 | 148 |
| N_Spneumoniae_D39 | Streptococcus pneumoniae D39 | 373153.27 | NC_008533 | fig\|373153.27.peg.643 | SPD_0565 | YP_816064.1 | 150 |
| C_Spneumoniae_D39 | Streptococcus pneumoniae D39 | 373153.27 | NC_008533 | fig\|373153.27.peg.644 | SPD_0566 | YP_816065.1 | 148 |
| Smitis_OT25 | Streptococcus mitis strain OT25 | 28037.212 | JYGP01000002 | fig\|28037.212.peg.409 | TZ90_00447 | KJQ68083.1 | 318 |
| Smutans_UA159 | Streptococcus mutans UA159 | 210007.7 | NC_004350 | fig\|210007.7.peg.1517 | SMU.1699c | NP_722028.1 | 310 |
| **MnmM_Name in FigS4A** | | | | | | | |
| Spneumoniae_ST556 | Streptococcus pneumoniae ST556 | 1130804.3 | CP003357 | fig\|1130804.3.peg.717 | MYY_0699 | AFC94360.1 | 185 |
| Spneumoniae_TIGR4 | Streptococcus pneumoniae TIGR4 | 170187.11 | NC_003028 | fig\|170187.11.peg.682 | SP_0652 | NP_345157.1 | 185 |
| Spneumoniae_R6 | Streptococcus pneumoniae R6 | 171101.6 | NC_003098 | fig\|171101.6.peg.637 | spr0571 | NP_358165.1 | 185 |
| Smutans_UA159 | Streptococcus mutans UA159 | 210007.7 | NC_004350 | fig\|210007.7.peg.1516 | SMU.1697c | NP_722027.1 | 181 |
| Smitis_OT25 | Streptococcus mitis strain OT25 | 28037.212 | JYGP01000002 | fig\|28037.212.peg.410 | TZ90_00448 | | 185 |
| Spneumoniae_D39 | Streptococcus pneumoniae D39 | 373153.27 | NC_008533 | fig\|373153.27.peg.645 | SPD_0567 | YP_816066.1 | 185 |

### Supplemental Figures

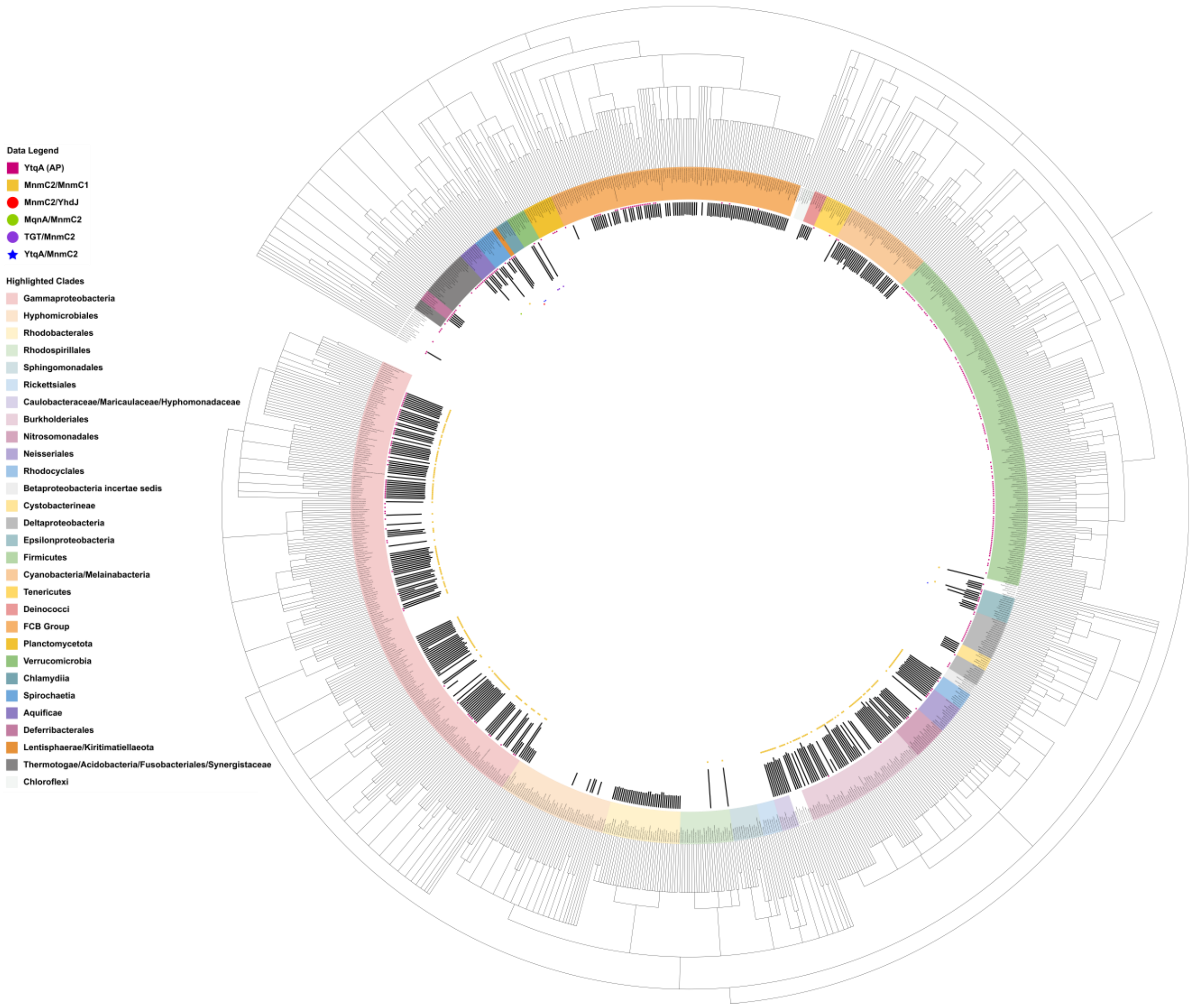

**Figure S1. Distribution of MnmC2 and MnmC2 fusions in the bacterial COG database genomes.** The presence of MnmC2 is represented by the respective homologs' lengths shown by the black histograms (2nd row, from outside-in) (**Supplemental data S3**). YtqA absence-presence is also shown by the fuchsia squares (1st row, outside-in). MnmC2/MnmC1 fusions are denoted by the goldenrod squares (3rd row). MnmC2/YhdJ fusions were noted by red circles (7th row) and MqnA(MqnD)/MnmC2 fusions by green circles (4th row), while the TGT/MnmC2 and YtqA/MnmC2 fusions were denoted by purple circles (5th row) and blue stars (6th row), respectively (derived from fusion data, **Supplemental data S3 and S4**). Clades of interest are highlighted in various colors across leaf labels, the key to which is provided left of the figure. An interactive version of this tree can be accessed at https://itol.embl.de/tree/8216522107187281668281889.

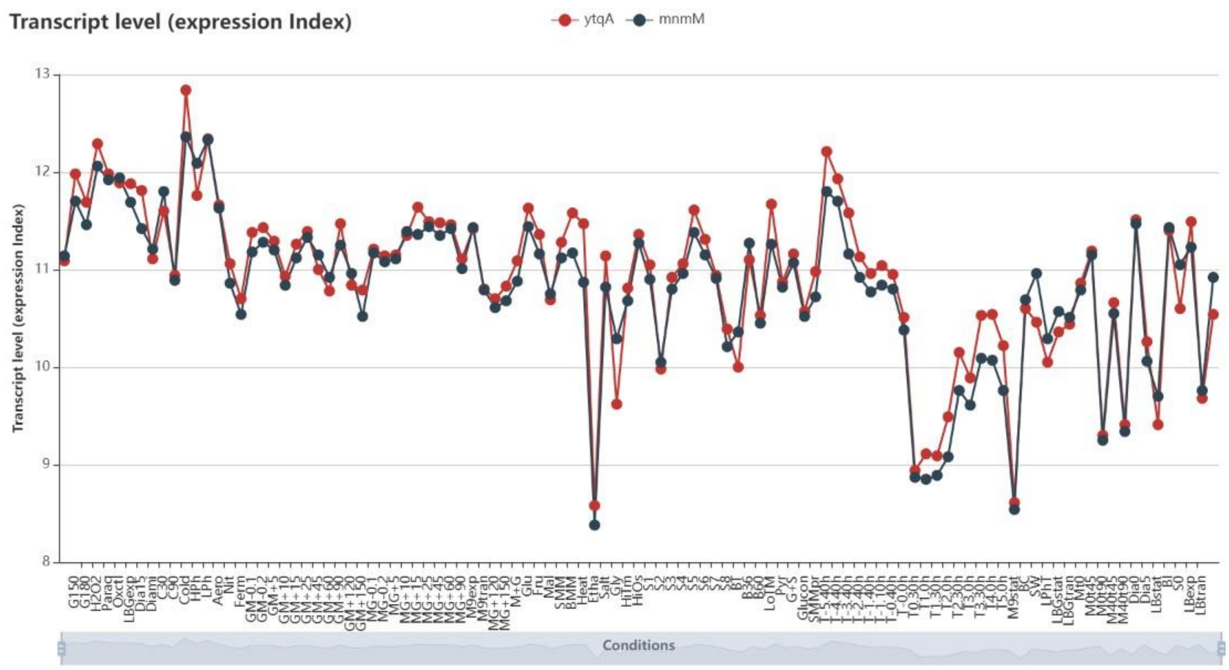

**Figure S2. Coexpression of YtqA and YtqB/MnmM in *B. subtilis.*** Data extracted from SubtiWiki http://subtiwiki.uni-goettingen.de/ expression browser using BSU_30480 (*ytqA*) and BSU_30490 (*mnmM*) as inputs.

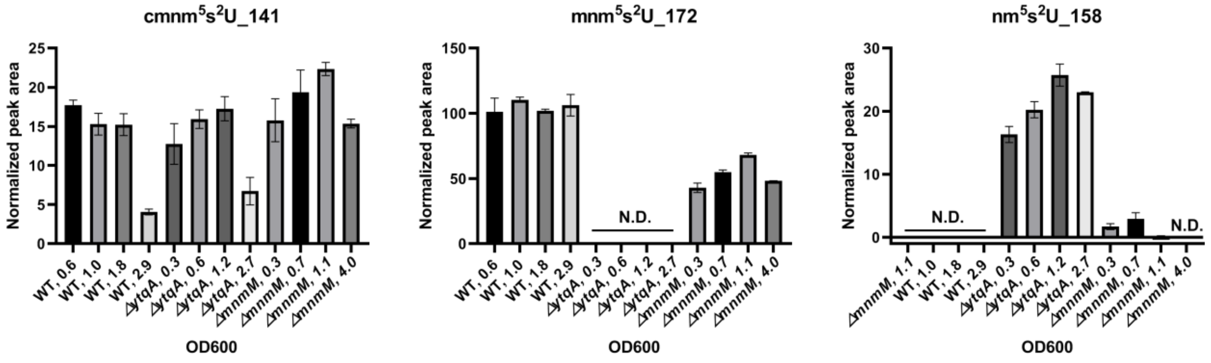

**Figure S3.** Analysis of $xm^5s^2U$ profiles in *Bacillus subtilis* WT and *ytqA* and *mnmM* mutant strains. Normalized peak area values for $cmnm^5s^2U$, $mnm^5s^2U$ and $nm^5s^2U$. The modification signals were based on specific transitions ($cmnm^5s^2U$_141, $mnm^5s^2U$_172 and $nm^5s^2U$_158) and were confirmed by both qualifier and quantifier transitions; 200 ng of hydrolysate was injected. tRNA samples were processed through VDCL2 method and analyzed through LC-MS method 2 as described in the methods section. N.D., Not detected. Samples were extracted at different stages of growth with OD ($A_{600nm}$) ranging from 0.3 to 4. Data represent the mean ±SD for 2 technical replicates.

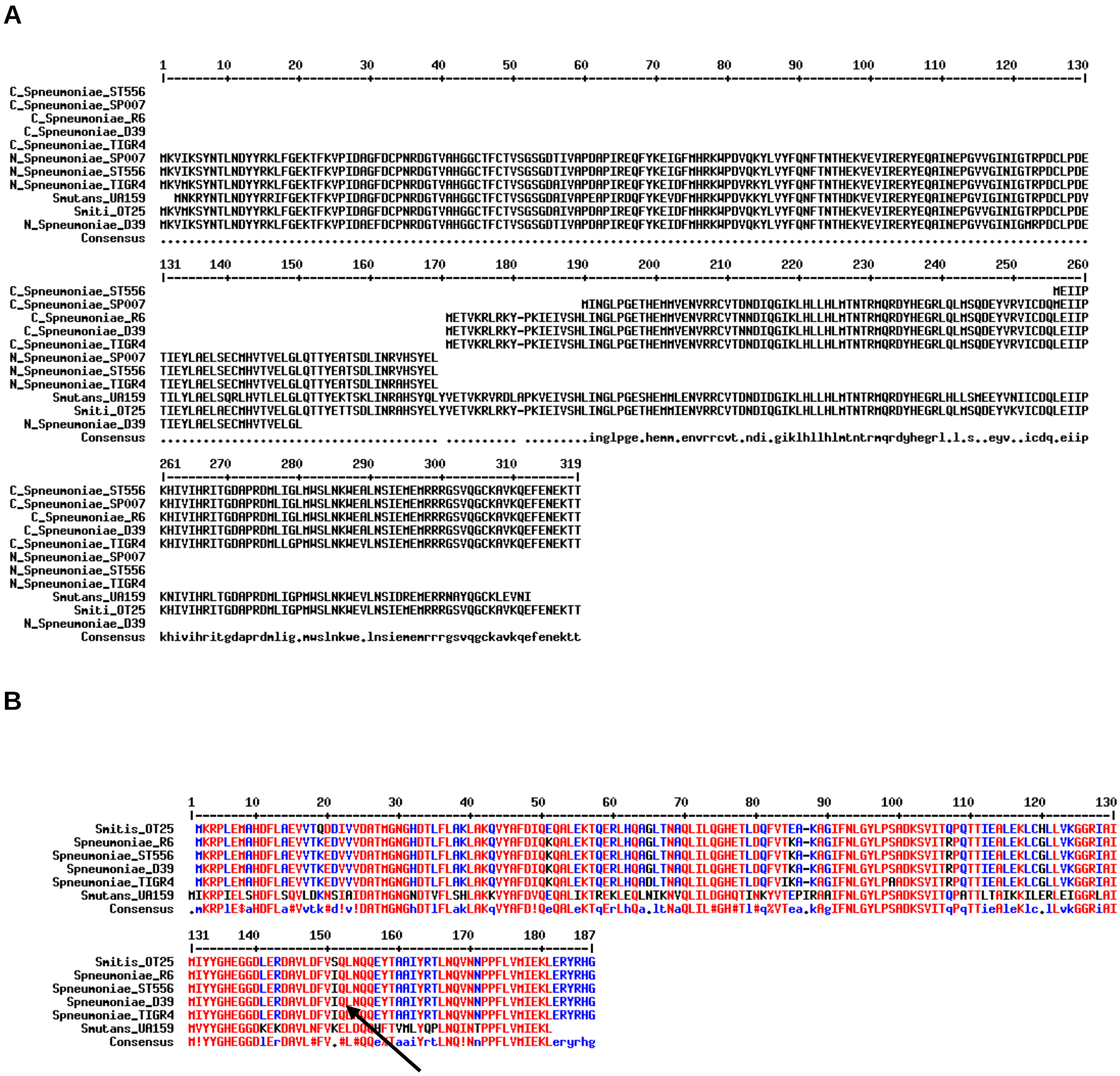

**Figure S4.** Sequence similarities of YtqA orthologs. (**A**) Alignments of truncated YtqA proteins from different *S. pneumoniae* strains (N-terminal and C-terminal halves are noted by a “C” or and an “N” in front of the species name) with full-length YtqA sequence from *S. mutans* (SMU_1699c) and *S. mitis* (TZ90_00447) using MultiAlin with default parameters. (**B**) Alignment of MnmM sequences from different *S. pneumoniae* strains with *S. mutans* (SMU_1697) and *S. mitis* (TZ90_00448) homologs. The location of the Q152STOP mutation that occurred in the IU1824 strain is shown with an arrow. Identifiers for all proteins in the alignments are listed in **Table S6**.

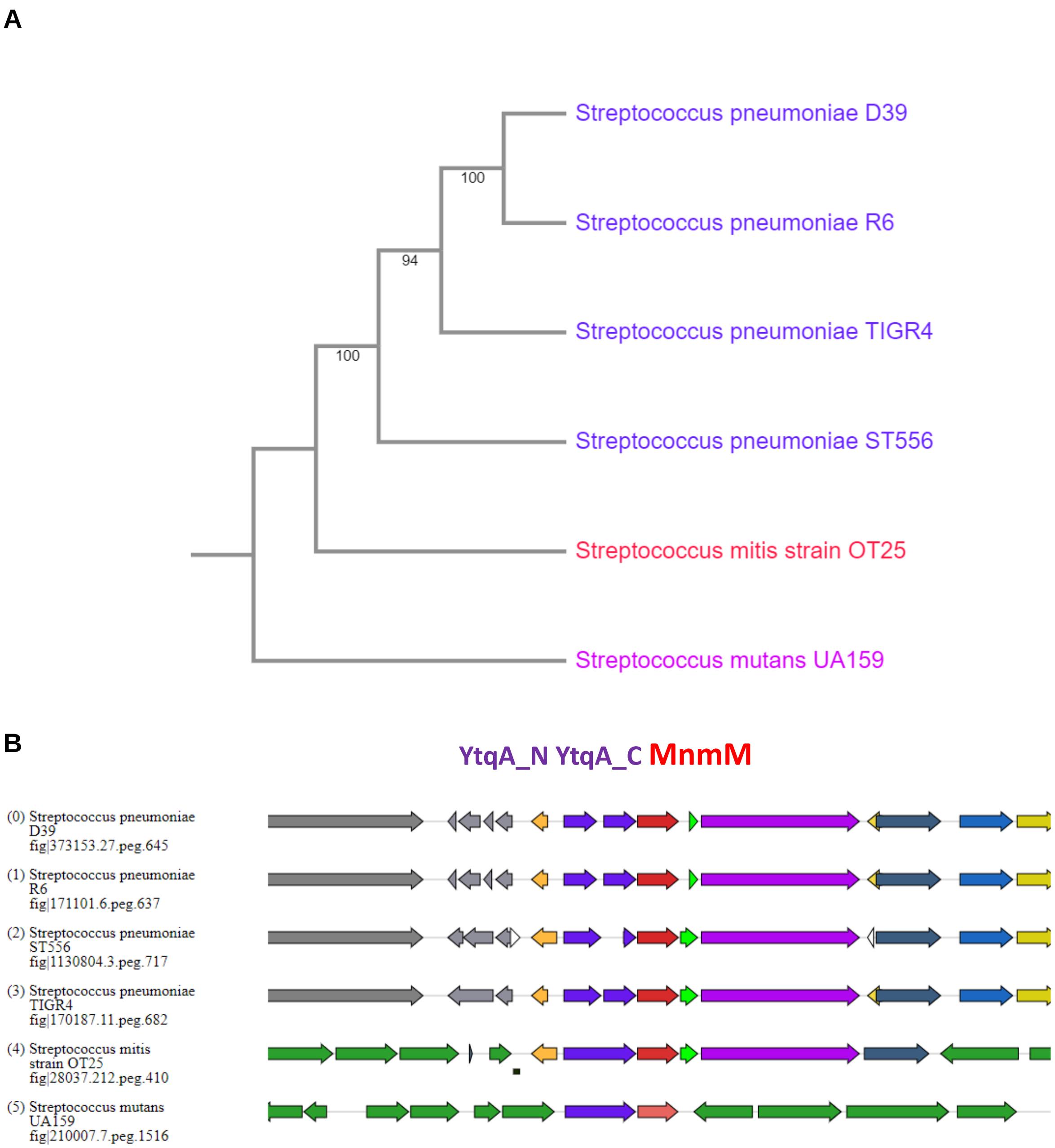

**Figure S5.** Diversity of *Streptococci* mnm$^5$s$^2$U biosynthetic genes. (**A**) The species tree shows the relationship between the *S. pneumoniae*, *S. mutans*, and *S. mitis* strains discussed in this study. The tree was built using BV-BRC bacterial Phylogenetic Tree Service that used the Codon Tree method that selects single-copy BV-BRC PGFams and analyzes aligned proteins and coding DNA from single-copy genes using the program RAxML (7). (**B**) Gene neighborhoods generated using Gizmogene around the *mnmM* ( in red) genes from the genomes shown in (A), showing the full-length or truncated ytqA (*ytqA_N* and *ytqA_C* genes ( in purple). Identifiers for all genomes and genes are given in **Table S6**.

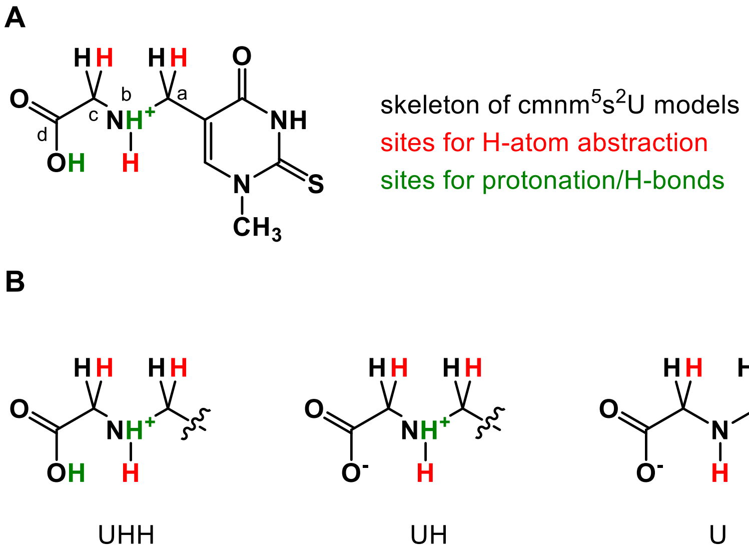

**Figure S6.** Models of cmnm$^5$s$^2$U employed in DFT calculations. Three different protonation states were considered (sites shown in green): UHH where the carboxylate and amide functional groups of the pendant glycine are protonated; UH in which only the amide is protonated; and U in which neither group is protonated. Three sites for H-atom abstraction were considered (sites shown in red): the glycine alpha carbon ($C_c$); the amide nitrogen ($N_b$); and the $^5$U-methylene carbon ($C_a$).

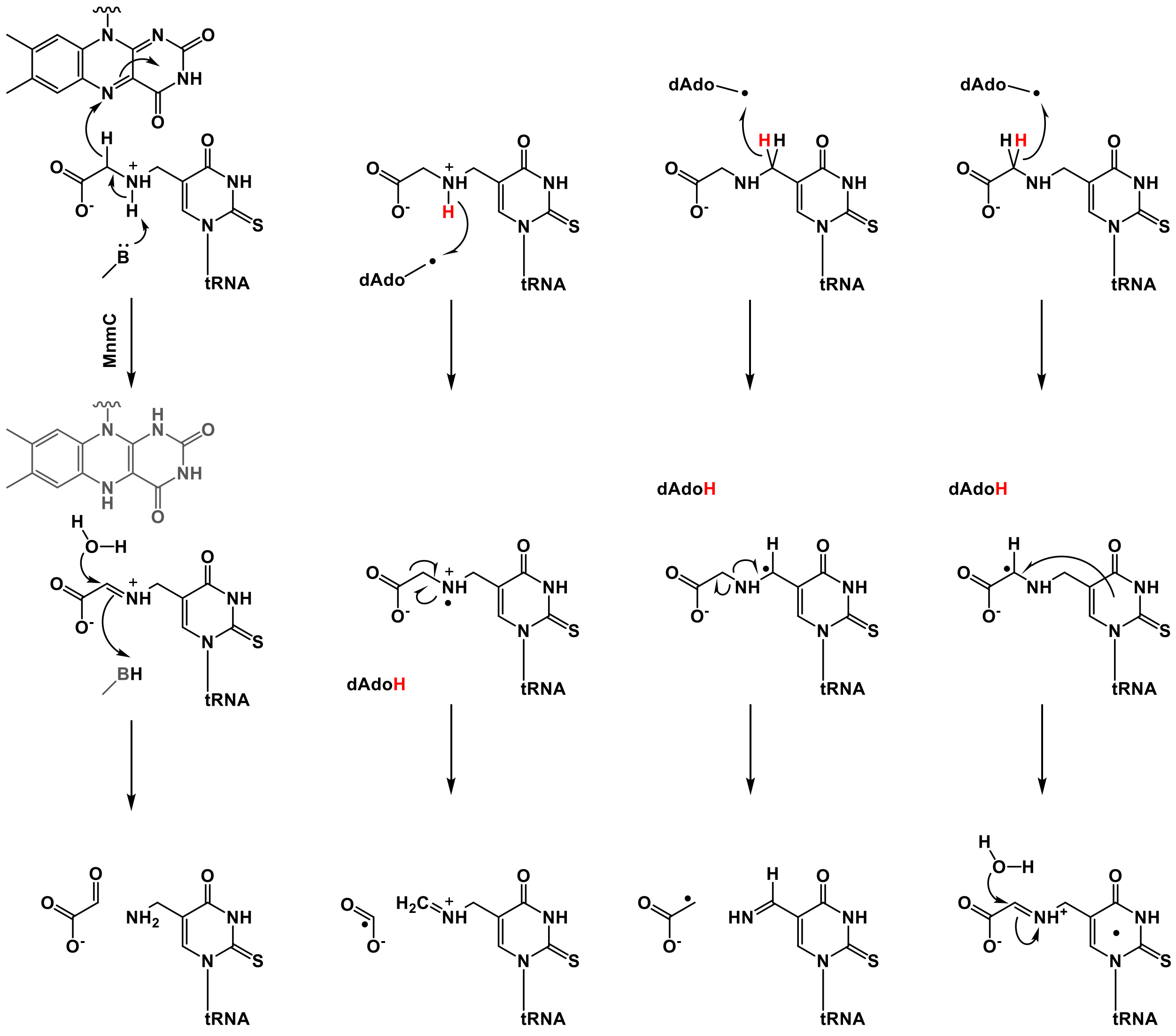

**Figure S7.** Hypothetical mechanisms for conversion of cmnm$^5$s$^2$U to nm$^5$s$^2$U.

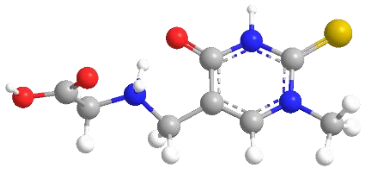

**UHH**

| atom | x (Å) | y (Å) | z (Å) |
|---|---|---|---|
| C(1) | 0.211 | 0.533 | -0.651 |
| C(2) | 0.335 | 1.318 | 0.570 |
| C(3) | 1.489 | 1.189 | 1.311 |
| O(4) | -0.769 | 0.611 | -1.431 |
| N(5) | 1.277 | -0.300 | -0.923 |
| C(6) | -0.859 | 2.135 | 0.996 |
| N(7) | 2.511 | 0.349 | 0.969 |
| C(8) | 3.717 | 0.264 | 1.809 |
| C(9) | 2.451 | -0.466 | -0.183 |
| S(10) | 3.643 | -1.509 | -0.640 |
| N(11) | -1.485 | 2.765 | -0.222 |
| C(12) | -2.847 | 3.342 | -0.073 |
| C(13) | -2.858 | 4.658 | -0.860 |
| O(14) | -4.077 | 5.177 | -0.946 |
| O(15) | -1.830 | 5.133 | -1.312 |
| H(16) | 1.641 | 1.758 | 2.242 |
| H(17) | 1.222 | -0.866 | -1.780 |
| H(18) | -0.601 | 2.919 | 1.734 |
| H(19) | -1.643 | 1.486 | 1.443 |
| H(20) | 4.609 | 0.540 | 1.214 |
| H(21) | 3.854 | -0.775 | 2.165 |
| H(22) | 3.600 | 0.951 | 2.666 |
| H(23) | -1.426 | 1.937 | -0.949 |
| H(24) | -0.901 | 3.549 | -0.608 |
| H(25) | -3.067 | 3.558 | 0.994 |
| H(26) | -3.621 | 2.642 | -0.443 |
| H(27) | -4.026 | 6.030 | -1.438 |

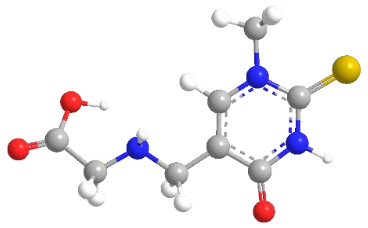

**UH**

| atom | x (Å) | y (Å) | z (Å) |
|---|---|---|---|
| C(1) | -0.059 | -0.800 | -0.077 |
| C(2) | -0.059 | 0.662 | 0.060 |
| C(3) | 1.121 | 1.292 | 0.349 |
| O(4) | -1.036 | -1.496 | -0.342 |
| N(5) | 1.214 | -1.376 | 0.127 |
| C(6) | -1.373 | 1.370 | -0.158 |
| N(7) | 2.323 | 0.633 | 0.532 |
| C(8) | 3.538 | 1.390 | 0.848 |
| C(9) | 2.415 | -0.756 | 0.423 |
| S(10) | 3.833 | -1.607 | 0.629 |
| N(11) | -1.374 | 2.749 | 0.345 |
| C(12) | -2.593 | 3.524 | 0.073 |
| C(13) | -2.264 | 5.027 | -0.079 |
| O(14) | -3.102 | 5.901 | -0.041 |
| O(15) | -0.952 | 5.240 | -0.298 |
| H(16) | 1.169 | 2.387 | 0.449 |
| H(17) | 1.264 | -2.397 | 0.040 |
| H(18) | -2.178 | 0.734 | 0.283 |
| H(19) | -1.595 | 1.402 | -1.249 |
| H(20) | 4.307 | 1.221 | 0.070 |
| H(21) | 3.961 | 1.047 | 1.813 |
| H(22) | 3.280 | 2.463 | 0.903 |
| H(23) | -0.567 | 4.314 | -0.238 |
| H(24) | -1.193 | 2.738 | 1.359 |
| H(25) | -3.401 | 3.410 | 0.828 |
| H(26) | -3.021 | 3.194 | -0.898 |

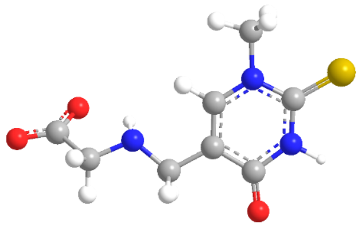

**U**

| atom | x (Å) | y (Å) | z (Å) |
|---|---|---|---|
| C(1) | -0.025 | -0.762 | -0.345 |
| C(2) | -0.109 | 0.648 | 0.029 |
| C(3) | 1.017 | 1.272 | 0.493 |
| O(4) | -0.929 | -1.481 | -0.770 |
| N(5) | 1.279 | -1.303 | -0.156 |
| C(6) | -1.413 | 1.405 | -0.071 |
| N(7) | 2.251 | 0.648 | 0.633 |
| C(8) | 3.400 | 1.409 | 1.122 |
| C(9) | 2.422 | -0.688 | 0.309 |
| S(10) | 3.881 | -1.511 | 0.459 |
| N(11) | -1.253 | 2.798 | 0.257 |
| C(12) | -2.466 | 3.609 | 0.331 |
| C(13) | -2.185 | 5.001 | -0.389 |
| O(14) | -3.031 | 5.902 | -0.203 |
| O(15) | -1.127 | 5.001 | -1.096 |
| H(16) | 0.937 | 2.328 | 0.800 |
| H(17) | 1.388 | -2.293 | -0.397 |
| H(18) | -2.136 | 0.920 | 0.629 |
| H(19) | -1.847 | 1.197 | -1.091 |
| H(20) | 4.218 | 1.392 | 0.374 |
| H(21) | 3.796 | 0.955 | 2.052 |
| H(22) | 3.074 | 2.448 | 1.308 |
| H(23) | -0.732 | 3.334 | -0.491 |
| H(24) | -2.802 | 3.801 | 1.376 |
| H(25) | -3.334 | 3.142 | -0.202 |

**Figure S8.** Geometry optimized models and *xyz* coordinates (Å) of cmnm$^5$s$^2$U that were further geometry-optimized after H-atom abstraction.

**Supplemental data description**

**Data S1**

List of the 968 organisms in the COG database that encode the three proteins MnmA, MnmG and MnmE  and distribution of the COG4121( MnmC) in those genomes.

**Data S2**

Summary of MnmC1/MnmC2/MnmC1C2 distributions across bacterial orders in the benchmark set of 968 genomes.

**Data S3**

Domain analysis of MnmC1/MnmC2/MnmC1C2 (via CDD Search) in homologs present in benchmark set of 968 genomes

**Data S4**

(a) List of YtqA/mmM, MnmC1/MnmM and YtqA/MnmC2 fusion proteins present in InterPro; (b) YtqA/YhcC family proteins found in the benchmark set of 968 genomes.

**Data S5**

YtqA/YhcC gene clusters (a) all, (b) those with mnmM. YtqAs taken from initial COG Db fusion analysis (unless Valerie wants me to go get these additional 40 UniProt fusion gene cluster data from KEGG); (c) YtqA/YhcC-physically clustered gene analyses identifying methylase/methyltransferase domains via three methods, results in numeric form (1, BLASTp; 2, CDD Search; 3, InterProScan); (d) YtqA-physically clustered gene analyses identifying methylase/methyltransferase domains via three methods, results with all hit sequence identifiers (1, BLASTp; 2, CDD Search; 3, InterProScan); (e) summary metrics for different methods determining methylase/methyltransferase domains among genes physically-clustered with YtqA/YhcC

**Data S6**

MnmC1/MnmC2/MnmC1C2, and physically clustered YtqA-MnmM occurrence data across bacterial orders. Data for each set of sequences includes the incidence counts across genomes and the calculated frequency for each bacterial order. These different sets of  data was used  to generate Fig. 2. All column counts/percentages were determined independently of each other, and, therein, MnmC1 and MnmC2 counts could be part of fusions or non-fused free-standing domains (*factually, however, the latter is not the case for MnmC1 as its occurrence was always observed in the form of a fusion).

**Data S7**

Differential YtqA gene cluster, MnmC1/MnmC2/MnmC1C2 fusion analysis data summary tables. For organisms containing YtqA from the benchmark genome set (filtered for positive MnmAGE status), the distributions of retrieved YtqA gene cluster data across genomes of select MnmC1/MnmC2/MnmC1C2 encoding statuses (subtables are labeled with their respective genome statuses).

**Data S8**

Complete YtqA/YhcC gene cluster, MnmC1/MnmC2/MnmC1C2 fusion analysis data mapping to SSN Figure: (a) "Node Count Cluster Number" mapped to "Sequence Count Cluster Number" for SSN mapping;"Node Count Cluster Number" is the actual cluster number within the SSN figure (follows visual order of appearance from top-left to bottom-right). (b) YtqA/YhcC gene cluster data mapped to an exported table of the AS100 repnode 55 YtqA/YhcC family SSN. (c) YtqA/YhcC gene cluster data-to-SSN cluster mapped data master sheet. (d) SSN-mapped YtqA/YhcC gene cluster data was transformed relative to unique metabolic category per gene cluster; data was used in subsequent metabolic category and gene symbol/family name analyses. (e) Taxonomic group distributions YtqA/YhcC gene clusters, MnmC1/MnmC2/MnmC1C2 fusion analysis data mapping to SSN. (f) Taxonomic group distributions YtqA/YhcC gene clusters, MnmC1/MnmC2/MnmC1C2 fusion analysis data mapping to SSN expanded relative to SSN cluster numbers.

**Data S9**

SSN file
