## SupplementalDataKey for "Alternate routes to mnm^5^s^2^U synthesis in Gram-positive bacteria"

Updated: CJR, Nov. 10, 2023

Data S1
MnmAGE COG absence-presence data; known members of the Mnm thiolation pathway, biosynthesis of mnm5s2U34

Data S2
MnmC1/MnmC2/MnmC1C2 distributions, general, across all COG bacterial genomes

Data S3
Domain analysis of MnmC1/MnmC2/MnmC1C2 (via CDD Search)

Data S4
(a) YtqA fusion analysis (InterPro); (b) YtqAs (all across MnmAGE bacterial genomes) (via COG Db)

Data S5
YtqA gene clusters (a) all, (b) those with mnmM. YtqAs taken from initial COG Db fusion analysis (unless Valerie wants me to go get these additional 40 UniProt fusion gene cluster data from KEGG); (c) YtqA-physically clustered gene analyses identifying methylase/methyltransferase domains via three methods, results in numeric form (1, BLASTp; 2, CDD Search; 3, InterProScan); (d) YtqA-physically clustered gene analyses identifying methylase/methyltransferase domains via three methods, results with all hit sequence identifiers (1, BLASTp; 2, CDD Search; 3, InterProScan); (e) summary metrics for different methods determining methylase/methyltransferase domains among genes physically-clustered with YtqA

Data S6
MnmC1/MnmC2/MnmC1C2, and physically clustered YtqA-MnmM occurrence data across bacterial orders. Data for each set of sequences includes the incidence counts across genomes and the calculated frequency for each bacterial order. These data would be mapped to a summary phylogenetic tree for the generation of Fig. 2. All column counts/percentages were determined independent of each other, and, therein, MnmC1 and MnmC2 counts could be part of fusions or non-fused free-standing domains (*factually, however, the latter is not the case for MnmC1 as its occurrence was always observed in the form of a fusion) 


Data S7
Differential YtqA gene cluster, MnmC1/MnmC2/MnmC1C2 fusion analysis data summary tables taxonomic group distributions relative to SSN cluster numbers


Data S8
Complete YtqA gene cluster, MnmC1/MnmC2/MnmC1C2 fusion analysis data mapping to SSN Figure: (a) "Node Count Cluster Number" mapped to "Sequence Count Cluster Number" for SSN mapping;"Node Count Cluster Number" is the actual cluster number within the SSN figure (follows visual order of appearance from top-left to bottom-right). (b) YtqA gene cluster data mapped to exported table of the AS100 repnode 55 YtqA family SSN. (c) YtqA gene cluster data-to-SSN cluster mapped data master sheet. (d) SSN-mapped YtqA gene cluster data transformed relative to unique metabolic category per gene cluster; data used in subsequent metabolic category and gene symbol/family name analyses. (e) Taxonomic group distributions YtqA gene cluster, MnmC1/MnmC2/MnmC1C2 fusion analysis data mapping to SSN. (f) Taxonomic group distributions YtqA gene cluster, MnmC1/MnmC2/MnmC1C2 fusion analysis data mapping to SSN expanded relative to SSN cluster numbers.


Data S9
SSN file
